# The chromatin remodeler Ino80 mediates alternative RNAPII pausing site determination

**DOI:** 10.1101/2021.04.02.438286

**Authors:** Youngseo Cheon, Sungwook Han, Taemook Kim, Daeyoup Lee

## Abstract

Promoter-proximal pausing of RNA polymerase II (RNAPII) is a critical step in early transcription elongation for the precise regulation of gene expression. Here, we provide evidence of promoter-proximal pausing-like distributions of RNAPII in *S. cerevisiae*. We found that genes bearing an alternative pausing site utilize Ino80p to properly localize RNAPII pausing at the first pausing site and to suppress the accumulation of RNAPII at the second pausing site, which is tightly associated with the +1 nucleosome. This alternative pausing site determination was dependent on the remodeling activity of Ino80p to modulate the +1 nucleosome position and might be controlled synergistically with Spt4p. Furthermore, we observed similar Ino80-dependent RNAPII pausing in mouse embryonic stem cells (mESCs). Based on our collective results, we hypothesize that the chromatin remodeler Ino80 plays a highly conserved role in regulating early RNAPII elongation to establish intact pausing.

## Introduction

Emerging evidence indicates that promoter-proximal pausing is a decisive step in transcription that supports the precise control of gene expression in metazoans (*1*). The establishment and release of paused RNAPII are strictly regulated by several factors during the early transcription elongation stage. Early biochemical studies using a purified system provided key mechanistic insights into pausing, showing that it is governed by two critical factors: DRB-sensitivity-inducing factor (DSIF; the heterodimeric Spt4/Spt5 complex) (*2*) and negative elongation factor (NELF) (*3*). Advances in genomic technology have enabled researchers to track the position of elongation complexes genome-wide by several methods, such as by capturing actively elongating RNAPII (*4–6*) or selecting RNAPII-associated RNAs (*7, 8*). The use of deep sequencing methods revealed that the loss of NELF reduced but did not completely abolish promoter-proximal pausing (*9, 10*), suggested that NELF acted to stabilize pausing rather than initiate it. Recent studies revealed that RNAPII pausing is found in species that lack NELF homologs, such as *C. elegans* (*11*) and *S. pombe* (*12*). Even *E. coli* RNA polymerases have been shown to pause at the start of the lambda gene (*13*). Further, the capture of nascent transcripts by native elongating transcript sequencing (NET-seq) in *S. cerevisiae* revealed that well-expressed genes exhibit a modest accumulation of read density downstream of the transcription start site (TSS) (*7*), which caused the non-uniform distribution of transcription elongation across genes. Overall, these previous studies have suggested that a conserved regulatory mechanism is involved in the early transcription elongation of yeast.

The nucleosome poses a strong barrier for RNAPII passage at various stages of transcription, and cells benefit from employing highly conserved chromatin remodelers to overcome these physical barriers (*14*). Several studies have shown that pausing occurs in close proximity to nucleosomes (*7, 12, 15, 16*). This suggested that nucleosomes can physically block the elongation of RNAPII, and that this collision causes RNAPII to pause. A genome-wide study targeting mouse Chd1 revealed that an ATPase inactive form of Chd1 results in a particular increase of RNAPII within the promoter regions (*17*), implying that chromatin remodeling could affect the promoter escape and subsequent pause-release of RNAPII. However, it remains unclear whether chromatin remodelers regulate nucleosome architecture to tune promoter-proximal pausing in the early elongation stage.

The chromatin remodeler, Ino80, has been shown to play a key role in the regulation of RNAPII at transcribed genes through its remodeling activity (*18*). The most well-known function of Ino80 is to exchange the highly conserved histone variant H2A.Z for H2A (*19, 20*). However, two recent studies disputed this function (*21, 22*). Several studies have suggested that Ino80p has the intrinsic capability of nucleosome spacing, and that it helps organize the intact nucleosome architecture around the promoter to regulate transcription (*23, 24*). Ino80 is largely enriched at the TSS of most genes in yeast and mammals (*20, 25*) and it has suggested that the recruitment of Ies6, the component of Ino80 complex, to the 5’ end of genes caused removal of histone H3-containing nucleosomes for gene expression in *S. pombe*. (*26*). Ino80 has also been shown to physically interact with the elongating RNAPII (*27, 28*). Nevertheless, the detailed mechanisms through which Ino80 regulates transcription elongation are currently unknown.

Here, we reveal the non-uniform distribution of elongating RNAPII in *S. cerevisiae* and show that it resembles promoter-proximal pausing in metazoans. Using the auxin-inducible degron (AID) system (*29, 30*), we found that Ino80p plays a key function in determining the position of RNAPII pausing in budding yeast. The genes whose pausing sites are regulated by Ino80p exhibited the use of an alternative pausing site rather than a focused pausing peak. The chromatin remodeling activity of Ino80p to properly localize the +1 nucleosome is essential to suppress RNAPII pausing at the second pausing site, thereby facilitating the utilization of the first pausing site. Further, we observed similar Ino80-dependent RNAPII pausing in mESCs, suggesting that Ino80 plays a highly conserved role in the regulation of RNAPII pausing.

## Results

### PRO-seq reveals genome-wide promoter-proximal pausing-like distributions in *S. cerevisiae*

To investigate the genome-wide distribution of elongation-competent RNAPII in *S. cerevisiae*, we first used Precision Run-On sequencing (PRO-seq) and Precision Run-On 5’ cap sequencing (PRO-cap) with 2-biotin run-on (biotion-11-CTP and UTP) (*6, 12*) (table S1). To more precisely define the transcription initiation sites, we chose the single base pair with the most PRO-cap reads within 250 bases upstream and downstream of the annotated TSS. We herein refer to the newly defined observed TSS as a “TSS” unless otherwise noted. Unexpectedly, PRO-seq revealed that transcription elongation was non-uniformly distributed: It was concentrated near TSSs, with a pattern resembling that associated with promoter-proximal pausing in metazoans (Fig. 1A; See black line of the profile). This was surprising because a previous study using PRO-seq in *S. cerevisiae* had captured a relatively uniform distribution of RNAPII across genes (*12*). To examine the significance of these apparent pausing-like features, we next classified genes as being paused or not paused, as described in the previous study (*12*). We identified 2,599 (45.5%) high-confidence paused and 2,099 (37.7%) not-paused genes among 5,315 filtered protein-coding genes (Fig. 1, A and B). The prevalence of pausing in *S. cerevisiae* was thus higher than that observed in *S. pombe* (28%) (*12*) and human (41%) (*4*) but lower than that observed in *D. melanogaster* (63%) (*9*). However, these differences in the relative number of paused genes could be a consequence of using different methods or gene sets to define RNAPII pausing (*31*).

**Fig. 1.**
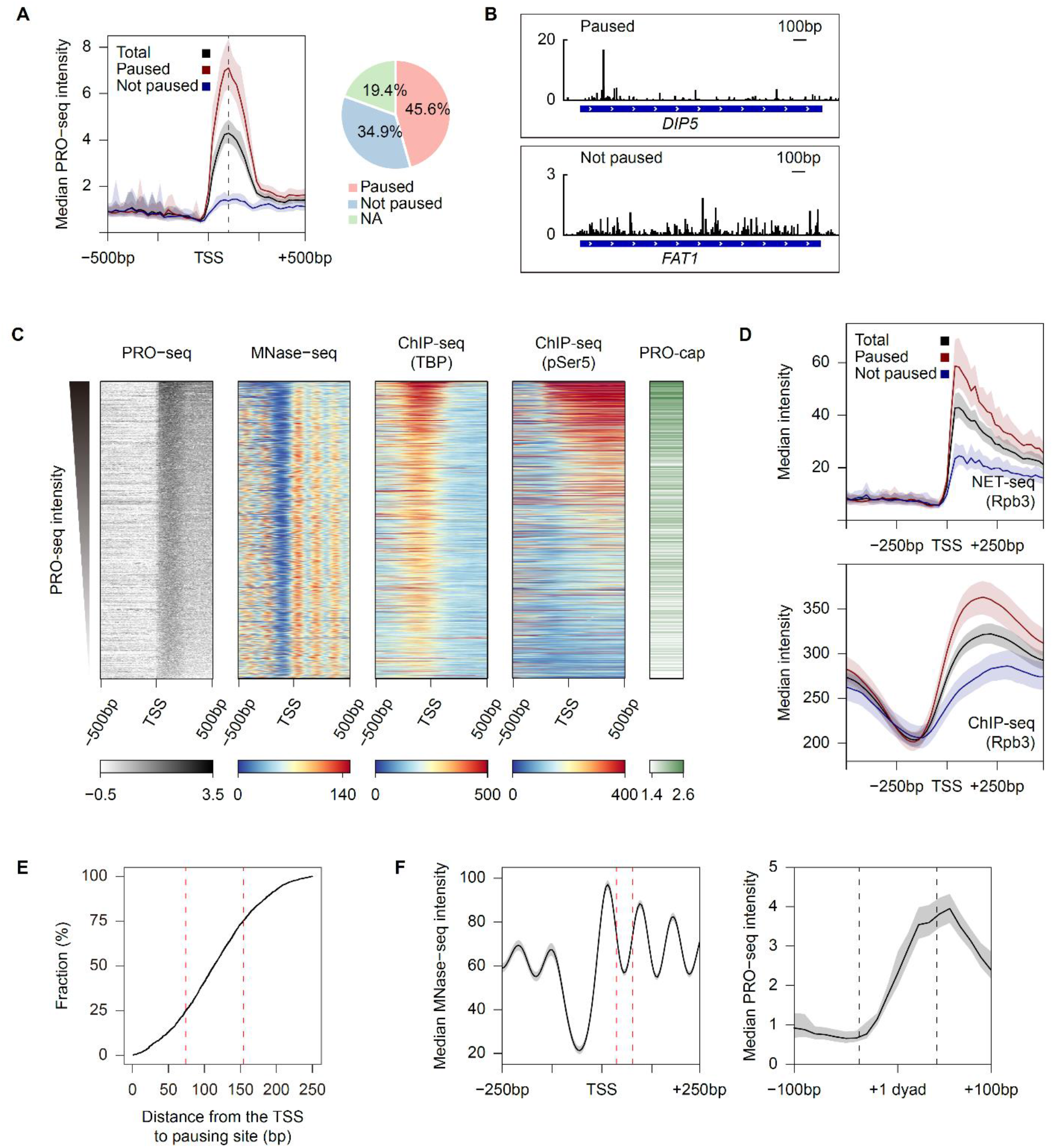
PRO-seq reveals a non-uniform distribution of transcription elongation genome-wide in *S. cerevisiae* resembling promoter-proximal pausing in metazoans. (**A**) Average profile showing the median PRO-seq intensities of paused and not paused genes. Pie chart indicates the percentage of paused (N = 2,599) and not paused (N = 1,990) genes among the total filtered protein-coding (N = 5,315) genes. NA indicate genes which were not classified as either being paused or not paused. (**B**) Genome browser view of PRO-seq signals for representative paused or not paused genes. (**C**) Heatmaps showing PRO-seq, PRO-cap, existing MNase-seq (GSM3304635), TBP ChIP-seq (GSM3452564) and pSer5 ChIP-seq (GSM3452562) signals. Genes were sorted by the PRO-seq signals at their promoter-proximal regions. The PRO-cap intensity reflected read counts within 250bp upstream and downstream of the TSS. (**D**) Average Rpb3 NET-seq (GSM617027) and ChIP-seq (GSM2813906) profiles centered on the TSS. Processed sequencing files downloaded from the NCBI Gene Expression Omnibus (GEO) were used for this analysis. (**E**) Cumulative curve analyzing the distance from the TSS to pausing site for paused genes. The red dotted lines represent the 25^th^ (74 bp) and 75^th^ (154 bp) percentiles. (**F**) Average profiles showing median MNase-seq intensity (GSM3304635) at the TSS (Left) and the median PRO-seq intensity at the +1 nucleosome dyad (Right). The +1 dyad was defined by the improved nucleosome-positioning algorithm, iNPS from an existing MNase-seq data (GSM3304635) (*63*). The two red dotted lines represent the 25^th^ (74 bp) and 75^th^ (154 bp) percentiles of the pausing site (Left) and the two black dotted lines represent the position of the +1 nucleosome (Right; 75 bp upstream and downstream of the +1 dyad). All PRO-seq data were generated from auxin-untreated Ino80-AID cells, and combined biological replicates were used. For average profiles, medians reflect either the 20-bp bin (PRO-seq) or the 10-bp bin (MNase-seq). For heatmaps, signals reflect the 10-bp bin around the indicated site. The PRO-seq, PRO-cap, and ChIP-seq intensities are presented in spike-in-normalized reads per million. MNase-seq data are presented in Gaussian smoothing-normalized reads per million.

We next sought to identify the general features of the paused genes in *S. cerevisiae*. To investigate whether the obtained PRO-seq reads were related to transcriptional activity or nucleosome density, we generated heatmaps of our data sets obtained from PRO-seq, PRO-cap, and existing data sets obtained using MNase-seq (*29*) and ChIP-seq against TBP and phosphor-Ser5 of RNAPII C-terminal domain (pSer5) (*32*). The RNAPII intensity within the promoter-proximal regions (TSS to TSS+250bp) generally correlated with the transcriptional activity (Fig. 1C); in this, our results were consistent with those of a previous study (*4*). In addition, the heatmaps showed that the higher the PRO-seq intensity, the lower the nucleosome occupancy and the wider the nucleosome free region (NFR) (Fig. 1C); this, too, was in accordance with earlier reports (*33*). Further, we compared our PRO-seq data with previously reported data sets obtained using Rpb3 NET-seq (*7*) and ChIP-seq (*34*). Consistent with our gene classification results, the NET-seq and ChIP-seq data displayed much higher enrichment at the TSS of paused genes compared to not-paused genes (Fig. 1D). Overall, we concluded that slowed RNAPII in *S. cerevisiae* is a conserved aspect of early transcription elongation that resembles promoter-proximal pausing in metazoans.

### Pausing-like feature in budding yeast is broader and more distal than that in metazoans

Despite this expectation, we observed a striking difference in the PRO-seq distributions of *S. cerevisiae* compared to metazoans. The peak of the PRO-seq enrichment within the promoter-proximal regions was located ∼100bp downstream of the TSS (Fig. 1A, Left). This localization was much farther downstream than the peak of the Global Run-On sequencing (GRO-seq) or the PRO-seq enrichment in metazoans, which showed the read peaks at ∼50bp downstream of the TSS in the sense strand (*4, 5*). Given that PRO-seq can track transcription elongation at almost single base-pair resolution, we defined the single-nucleotide of the maximum PRO-seq read within the promoter-proximal regions as the pausing site, as described in a recent study (*16*). The cumulative curve demonstrated that the 25^th^ and 75^th^ percentiles of the distance from the TSS to pausing site were 74 and 154 bp, respectively (Fig. 1E; The red dotted lines). To determine the association between pausing and the +1 nucleosome, we generated an average profile of existing MNase-seq data used in Fig. 1C around the TSS (Fig. 1F, Left; The 25^th^ and 75^th^ percentiles of the distance from the TSS to the pausing site are represented as the two red dotted lines) and our PRO-seq data around the +1 dyad, which was determined by the same MNase-seq data (Fig. 1F, Right). Interestingly, the majority of pausing sites were found to be located downstream of the +1 nucleosome. This is consistent with the previous report using NET-seq, which showed the peak of mean pause density downstream of the +1 nucleosome dyad (*7*). In contrast, RNAPII is generally paused upstream of the +1 dyad (*5, 15, 16*) in metazoans. This difference suggests that RNAPII undergoes a longer elongation period before pausing in budding yeast compared to metazoans and it could likely be attributed to differences in promoter structure. The +1 nucleosome typically includes a TSS for most yeast genes, whereas the +1 nucleosome of metazoans is located downstream of the TSS (*35, 36*).

### Loss of Ino80p causes variation in fitness and the PRO-seq pattern

Given the apparent role of Ino80p in transcription elongation (*18*), we used PRO-seq to investigate the role of Ino80p in nascent transcription at nearly single-nucleotide resolution. We first set out to map transcription elongation in wild-type and *ino80*Δ cells. Interestingly, *ino80*Δ cells exhibited a 5’-direction skew of the promoter-proximal peak in replicates 2 (*ino80*Δ_2) and 3 (*ino80*Δ_3) relative to that in wild-type cells (fig. S1A). However, these results were not comparable, as replicate 1 (*ino80*Δ_1) did not show a skewed PRO-seq distribution (fig. S1A). Consistent with our PRO-seq results, the cells of same batch used to generate *ino80*Δ_1 exhibited better growth than *ino80*Δ_2 and *ino80*Δ_3 (fig. S1B). These variations in the PRO-seq distribution and cellular fitness of *ino80*Δ cells seem to be attributed to the appearance of revertant due to the severe growth defect of *ino80*Δ. Thus, we employed an auxin-inducible degradation system (*29, 30*), with the goal of generating highly reproducible data and exclude the indirect effect of Ino80p in transcription elongation. Briefly, Ino80-AID strains were grown to mid-log phase in yeast peptone dextrose (YPD) containing ethanol (Ctrl). Ethanol was washed from the media and the cells were incubated with auxin (0.5mM) for 3 hrs (KD). Auxin was washed from the media and cells were incubated in auxin-free medium for an additional 3 hrs (Rescue) (fig. S1C). Western blot analysis confirmed the conditional depletion and recovery of Ino80p in an AID-tag-dependent manner after the 3 hrs incubations with or without auxin (fig. S1D).

### Ino80p is critical to the proper positioning of RNAPII pausing at the genes bearing an alternative pausing site

We carried out PRO-seq experiments to determine whether Ino80p knockdown (Ino80p-KD) caused a similar skewed pattern of the promoter-proximal peak in *ino80*Δ cells. The average profile of the median PRO-seq intensity for paused genes (N = 2,599) displayed a general skew of the promoter-proximal signal in the 5’ direction upon Ino80p-KD (fig. S2A). The pausing sites observed after 0 hr and 3 hrs of auxin treatment in Ino80-AID cells, which were assigned as described above, were designated “the 1^st^ pausing site” and “the 2^nd^ pausing site”, respectively. Cumulative curves and boxplots of the distance between the TSS and the pausing site revealed that the pausing site was significantly shifted in the 5’ direction upon Ino80p-KD (fig. S2B). To analyze this RNAPII transition in detail throughout the genome, we generated a heatmap of the PRO-seq log_2_ fold change around the 1^st^ pausing site. Interestingly, 12.6% of the paused genes (N = 2,599) were shifted toward 3’ upon Ino80p-KD; however, consistent with our cumulative curve and boxplot data, the 5’ shift was stronger and more frequent (28.5%) (fig. S2C). We chose transcripts that showed shifts of more than 30bp in their pausing site upon Ino80p-KD (to enable us to observe a clear change in position) and investigated whether the pausing site tended to be restored upon Ino80p rescue. Indeed, a boxplot analysis indicated that the changes of pausing site were significantly recovered under auxin withdrawal for both shifted gene sets (fig. S2D). In contrast, Ino80p rescue did not affect pausing sites in genes that were not shifted upon Ino80p-KD. These results suggest that Ino80p plays a previously unrecognized function in proper localization of RNAPII pausing sites in the early elongation stage for a subset of genes.

To select genes directly regulated by Ino80p, we chose transcripts that were shifted more than 30bp upstream in knockdown cells and restored in rescued cells (N = 221) and those that failed to show any shift under knockdown or subsequent restoration (N = 1,211). We referred to the former gene set as “shift-to-5’ genes” and the latter as “no-shift genes”. To examine whether the assigned pausing site precisely reflects the peak of pausing, we generated the average profile of the median PRO-seq signal around the TSS for shift-to-5’ genes. We observed that the PRO-seq peak exhibited striking upstream movement upon Ino80p-KD (Fig. 2A; The arrows and below dotted lines represent the median of the pausing sites). As expected, this shifted pausing was almost completely restored to the 1^st^ pausing site when Ino80p was rescued (Fig. 2A). Thus, our results suggest that this pausing site determination is totally Ino80p expression-dependent.

**Fig. 2.**
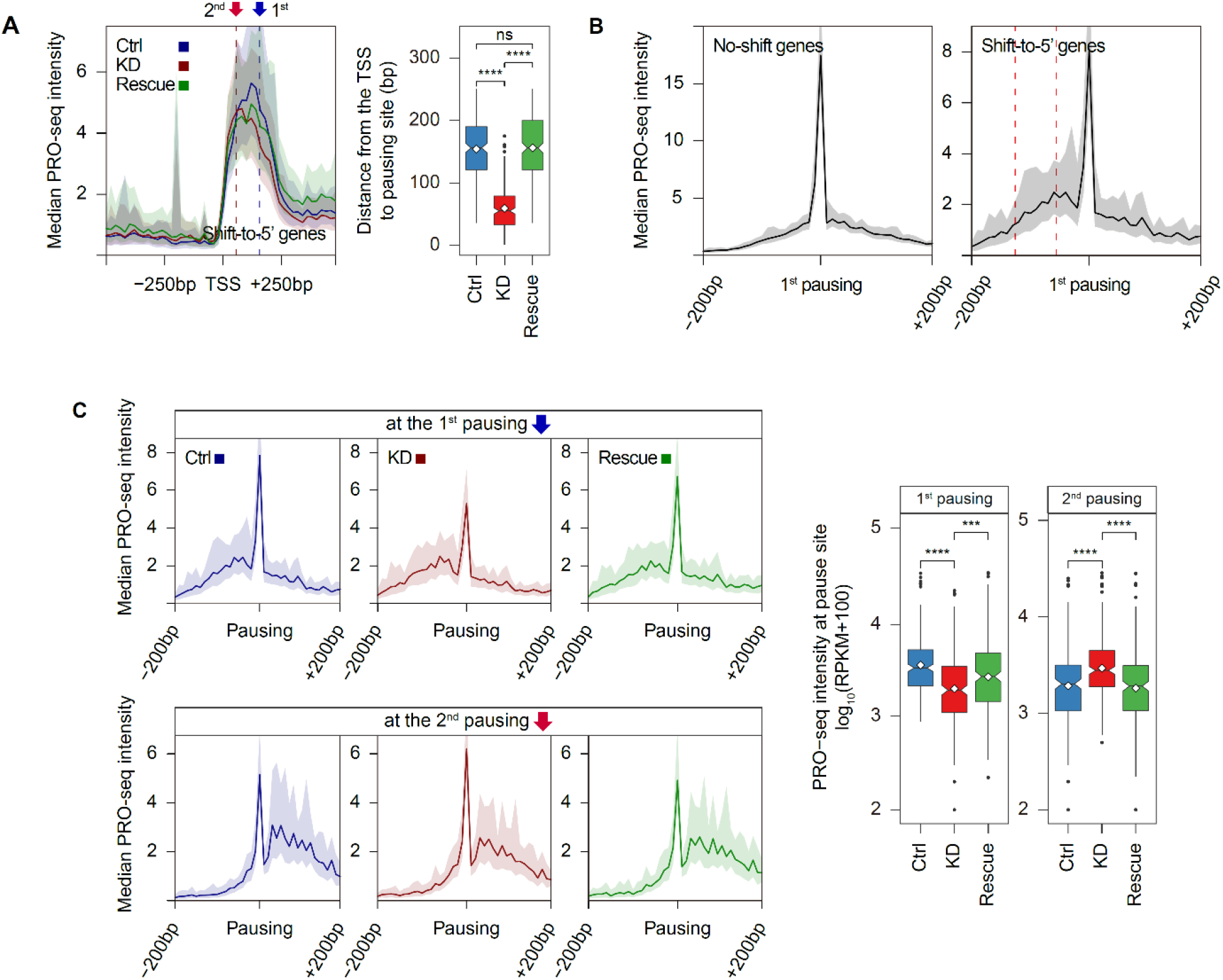
Pausing-site determination by Ino80p occurs at genes with alternative pausing sites. (**A**) Average profile indicating median PRO-seq intensities in Ino80-AID cells treated with auxin for 0 hr (Ctrl) and 3 hrs (KD) and rescued for 3 hrs after auxin removal (Rescue) for shift-to-5’ genes (N = 221). The arrows and below dotted lines represent the median of the 1^st^ (blue, 55bp) and 2^nd^ (red, 156bp) pausing sites. Medians reflect 20-bp bins. Boxplot shows the distance between the TSS and pausing site. (**B**) Average profiles representing median PRO-seq intensities in the control sample for either no-shift genes (N = 1,211) or shift-to-5’ genes (N = 221). The two red dotted lines (Right) represent the 25^th^ (-56 bp) and 75^th^ (-126 bp) percentiles of the 2^nd^ pausing site relative to the 1^st^ pausing site. Medians reflect 10-bp bins. (**C**) Average profiles displaying median PRO-seq intensities around either the 1^st^ or the 2^nd^ pausing site. Medians reflect 10-bp bins. Boxplot shows the PRO-seq intensity at the indicated single nucleotide in log_10_(RPKM+100). All data was generated using combined biological replicates. PRO-seq intensity was calculated using spike-in-normalized reads per million. Asterisks represent statistically significant differences, as calculated using the Wilcoxon test.

To investigate the general features of Ino80p-dependent genes, we compared the PRO-seq pattern at the 1^st^ pausing site under untreated conditions for no-shift genes and shift-to-5’ genes. Surprisingly, we observed only a small accumulation of PRO-seq signal upstream of the peak at the 1^st^ pausing site for shift-to-5’ genes, whereas no-shift genes displayed only a single sharp and distinct peak (Fig. 2B). The 25^th^ and 75^th^ percentiles of the 2^nd^ pausing site relative to the 1^st^ pausing site for shift-to-5’ genes (represented by the red dotted lines in the Fig. 2B) indicated that Ino80p-KD displaces RNAPII solely at locations where pausing can also occur but not a main pausing site. These results implied that Ino80p-dependent genes bear an alternative pausing site and utilize Ino80p to facilitate RNAPII pausing at the main pausing site. Further, we found that pausing at the 1^st^ pausing site was decreased upon Ino80p-KD and recovered upon Ino80p rescue (Fig. 2C, Upper) for shift-to-5’ genes, whereas pausing at the 2^nd^ pausing site showed an opposite tendency (Fig. 2C, Bottom). Thus, we propose that Ino80p is critical for the passage of RNAPII from the 2^nd^ pausing site to the 1^st^ pausing site for the establishment of intact pausing. We also noted that the elongating RNAPII at the 2^nd^ pausing site in untreated Ino80-AID cells was above the basal level (Fig. 2C, Bottom; Ctrl sample), which is consistent with the hypothesis that RNAPII could also pause at the 2^nd^ pausing site in the physiological state, but it does not represent a major pausing site. Supporting this idea, the sequence preferences at the 1^st^ and the 2^nd^ pausing site generated by WebLogo (*37*) exhibited a marked similarity (fig. S2E), suggesting the involvement of *cis*-acting nucleic acid sequences in RNAPII pausing. Overall, these results implied that Ino80p might play a pivotal role in determining proper pausing sites on genes bearing alternative pausing sites.

### The transition of the pausing site in Ino80p-KD is independent of both TSS usage and H2A.Z^Htz1^

We next determined whether the observed transition of RNAPII pausing was due to a defect in TSS usage. The precise transcription initiation sites for shift-to-5’ genes upon Ino80p-KD were identified by the maximum PRO-cap read within 100bp around the defined TSS in control samples. Histograms of the distance between the TSS in control and knockdown samples indicated that the majority of these genes exhibited no differences in transcription initiation site (fig. S2F, Left). This suggested that the Ino80p-dependent positioning of RNAPII pausing may result from a defect in transcription elongation rather than a defect in TSS usage. To further investigate whether the PRO-cap intensity was altered upon Ino80p-KD, we measured the log_2_ fold change of the PRO-cap signal 100bp around the TSS for shift-to-5’ or no-shift genes (fig. S2F, Right). Our boxplot analysis demonstrated that the changes in PRO-cap signal for shift-to-5’ genes showed a greater decrease than those for no-shift genes, suggesting that there may be an association between pausing site determination and the abundance of 5’ capped RNA.

Given that previous reports proposed a connection between Ino80p and H2A.Z^Htz1^ (*19, 22*), we questioned whether the transition of RNAPII could be caused by the insufficient removal of H2A.Z^Htz1^. To test this possibility, we carried out PRO-seq experiments in *htz1*Δ cells. The average profile, however, revealed that *htz1*Δ did not result in a skewed pattern of the promoter-proximal peak similar to that seen for the shift -to-5’ genes upon Ino80p-KD (fig. S2G, Left). Moreover, the distance of pausing site shift in Ino80p-KD showed no significant correlation with H2A.Z^Htz1^ enrichment in the +1 nucleosome, which is calculated from an existing MNase-ChIP-seq (*38*) (fig. S2G, Right). Thus, we conclude that the function of Ino80p in pausing site positioning is independent of its role in restricting the localization of H2A.Z^Htz1^, at least for shift-to-5’ genes. Consistent with this proposal, previous studies showed that the occupancy of H2A.Z^Htz1^ on chromatin is not altered under Ino80 depletion (*21, 22*).

### The transition of RNAPII pausing is closely associated with the +1 nucleosome

We questioned whether the transition of RNAPII upon Ino80p-KD was associated with the nucleosome architecture around the pausing site. To address this, we first analyzed the average profile of an existing MNase-seq data generated using untreated Ino80-AID cells (*29*), which is the same cell background used in our PRO-seq. Surprisingly, the nucleosome distribution relative to the 1^st^ pausing site of shift-to-5’ genes displayed a much better phase than that of no-shift genes (Fig. 3A). Also, the +1 nucleosome tended to be located in closer proximity to the 2^nd^ pausing site than the 1^st^ pausing site (Fig. 3A; The 25^th^ and 75^th^ percentiles of the 2^nd^ pausing site relative to the 1^st^ pausing site are represented as the two dotted lines). This implicated that the ability of Ino80p to suppress RNAPII pausing at the 2^nd^ pausing site in a nucleosome context-dependent manner. We also examined existing MNase-seq data obtained upon Ino80p-KD (*29*). While the nucleosome distribution for no-shift genes showed almost no difference, that for shift-to-5’ genes exhibited a moderate disturbance in nucleosome positioning and a decrease in the +1 nucleosome occupancy upon knockdown (Fig. 3B). When we performed the same analysis for MNase-seq data distributed by other laboratories (*32*), which differed slightly in the cell background, incubation temperature, and auxin treatment time, we obtained similar results (fig. S3, A and B).

**Fig. 3.**
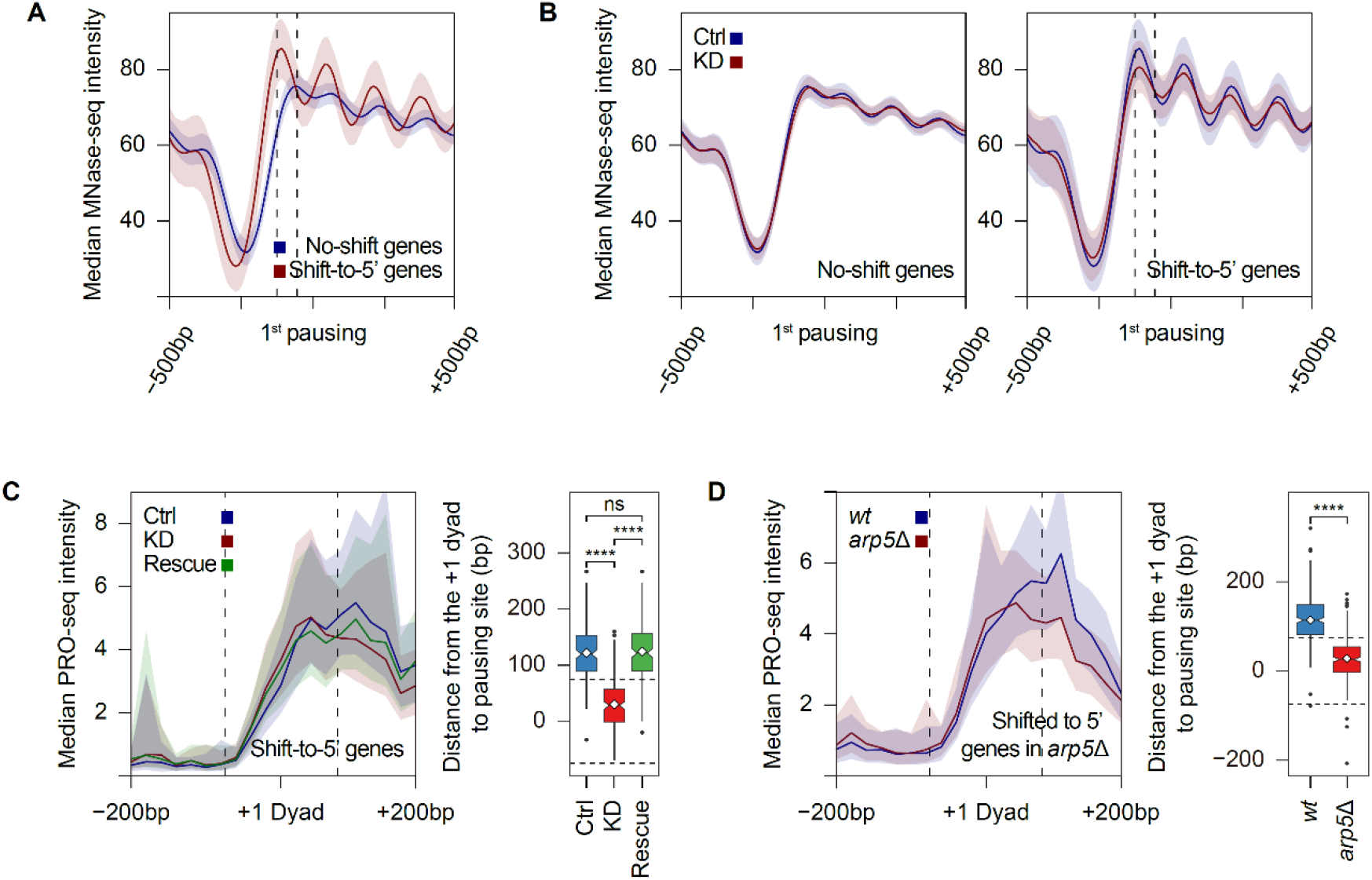
Chromatin remodeling activity of Ino80p in nucleosome positioning around the 1^st^ pausing site is critical for proper localization of pausing. (**A**) Average profiles of median MNase-seq intensities in control samples (GSM3304635) for no-shift genes (N = 1,211) and shift-to-5’ genes (N = 221) centered on the 1^st^ pausing site. (**B**) Average profiles of median MNase-seq intensities in control sample (GSM3304635) and Ino80p-KD sample (GSM3304637) for no-shift genes (Left) and shift-to-5’ genes (Right) at the 1^st^ pausing site. For Fig. 3, A and B, the two dotted lines indicate the 25^th^ (-56 bp) and 75^th^ (-126 bp) percentiles of the 2^nd^ pausing site relative to the 1^st^ pausing site. (**C** and **D**) Average profiles of median PRO-seq intensity for the indicated samples around the +1 dyad (defined in Fig. 1F) and boxplots of the distance from the +1 dyad to pausing site. Note that only nucleosomes overlapped with H3K4me3 ChIP-seq enrichment (GSM2507874) were used, in an effort to exclude false-positive nucleosomes. For (**C**), shift-to-5’ genes upon Ino80p-KD (N = 160) were used for analyses. For (**D**), pausing site shifted toward 5’ genes in *arp5*Δ (N = 264) were used for analyses. The two dotted lines in profiles and boxplots represented the position of the +1 nucleosome (75 bp upstream and downstream of +1 dyad). All PRO-seq data were generated using combined biological replicates. PRO-seq and MNase-seq intensity were calculated using spike-in-normalized reads per million and Gaussian smoothing-normalized reads per million, respectively. For average profiles, medians reflect either the 20-bp bin (PRO-seq) or the 10-bp bin (MNase-seq). Asterisks represented statistically significant differences, as calculated using the Wilcoxon test.

To continue addressing the association of the 2^nd^ pausing site with the +1 nucleosome, we next evaluated the PRO-seq distribution around the +1 dyad (defined in Fig. 1F). To discard false-positive nucleosome positions, we excluded nucleosomes that did not overlap the H3K4me3 ChIP-seq enrichment calculated from the existing data (*34*), as previously reported (*16*). We generated a composite profile centered on the +1 dyad and a boxplot of the distance to pausing site from the +1 dyad. No-shift genes were divided by whether they exhibited a pausing site outside (N = 463) or inside (N = 401) of the +1 nucleosome, to clearly distinguish the changes in PRO-seq distribution. Neither group of no-shift genes displayed any distinct change in pausing site relative to the +1 dyad upon Ino80p-KD (fig. S3C). In striking contrast, we observed that the transition from the 1^st^ to the 2^nd^ pausing site occurred through the +1 nucleosome for shift-to-5’ genes (N = 160) (Fig. 3C). It seemed that the large fraction of elongating RNAPII could not pass the +1 nucleosome upon Ino80p-KD. Rescue of Ino80p expression caused almost perfect restoration of the pausing site to the downstream of the +1 nucleosome, further supporting the direct function of Ino80p. Based on these findings, we propose that the regulation of the nucleosome positioning around the 1^st^ pausing site by Ino80p is critical to suppress RNAPII accumulation at the 2^nd^ pausing site.

### Ino80 remodeling activity might be critical in pausing site determination

To distinguish whether the regulation of nucleosome positioning around the pausing site depends on direct remodeling activity of Ino80p or other *trans*-activating factors associated with Ino80p, we carried out PRO-seq experiments in *arp5*Δ cells, which lack a component that is essential for the chromatin remodeling activity of Ino80p complex in *S. cerevisiae* (*39–41*). When we analyzed the defined paused genes as described for Ino80p-KD cells, we observed that a similar shift of pausing site toward the inside of the +1 nucleosome occurred solely for pausing site shifted toward 5’ genes (> 30bp) in *arp5*Δ (N = 264) (Fig. 3D and fig. S3D). Corroborating this, a Venn diagram analysis revealed that there was a significant overlap (*P* value = 1.52×10^-12^) between shift-to-5’ genes upon Ino80p-KD and genes showing an upstream shift of RNAPII pausing in *arp5*Δ (fig. S3E). Thus, we concluded that the chromatin remodeling activity of the Ino80p complex to the +1 nucleosome plays an important role in well-positioning of RNAPII pausing.

### The conserved pausing factor, Spt4p, plays a similar role in regulating alternative pausing site

Since a previous study suggested that the conserved pausing factor, Spt4p, is required to facilitate productive transcription elongation in *S. cerevisiae* (*12*), we postulated Spt4p is also critical for transitiong the elongating RNAPII from the 2^nd^ to the 1^st^ pausing site. To examine this, we conducted PRO-seq in *spt4*Δ cells. Using the same set of analyses performed with the PRO-seq data in Ino80p-KD, we selected pausing site shifted toward 5’ direction (> 30bp) gene sets in *spt4*Δ (N = 784). As expected, we observed a similar upstream skewed pattern of the PRO-seq peak in *spt4*Δ relative to wild-type within the promoter-proximal regions (Fig. 4, A and B). Further, PRO-seq signal in *spt4*Δ showed a prominent increase immediately downstream of the TSS, which was consistent with the previous report (*12*). There was a significant overlap (*P* value = 5.40×10^-8^) between shift-to-5’ genes upon Ino80p-KD and Spt4p-regulated genes (Fig. 4C).We also observed strong negative genetic interactions between *INO80* and both *SPT4* and *PAF1* (Fig. 4D), the latter of which has been shown to physically interact with DSIF and RNAPII and facilitate RNAPII pausing (*42–44*). Recent work showed that Spt4, Spt5, and the additional elongation factor Elf1 could cooperatively prevent RNAPII pausing at SHL(-6), SHL(-5) and SHL(-2) *in vitro* (*45*), suggesting that Spt4 plays a role in overcoming the nucleosome barrier. Based on these results, we propose that Ino80p and Spt4p synergistically regulate the ability of elongating RNAPII to travel along the +1 nucleosome and establish intact pausing in *S. cerevisiae*.

**Fig. 4.**
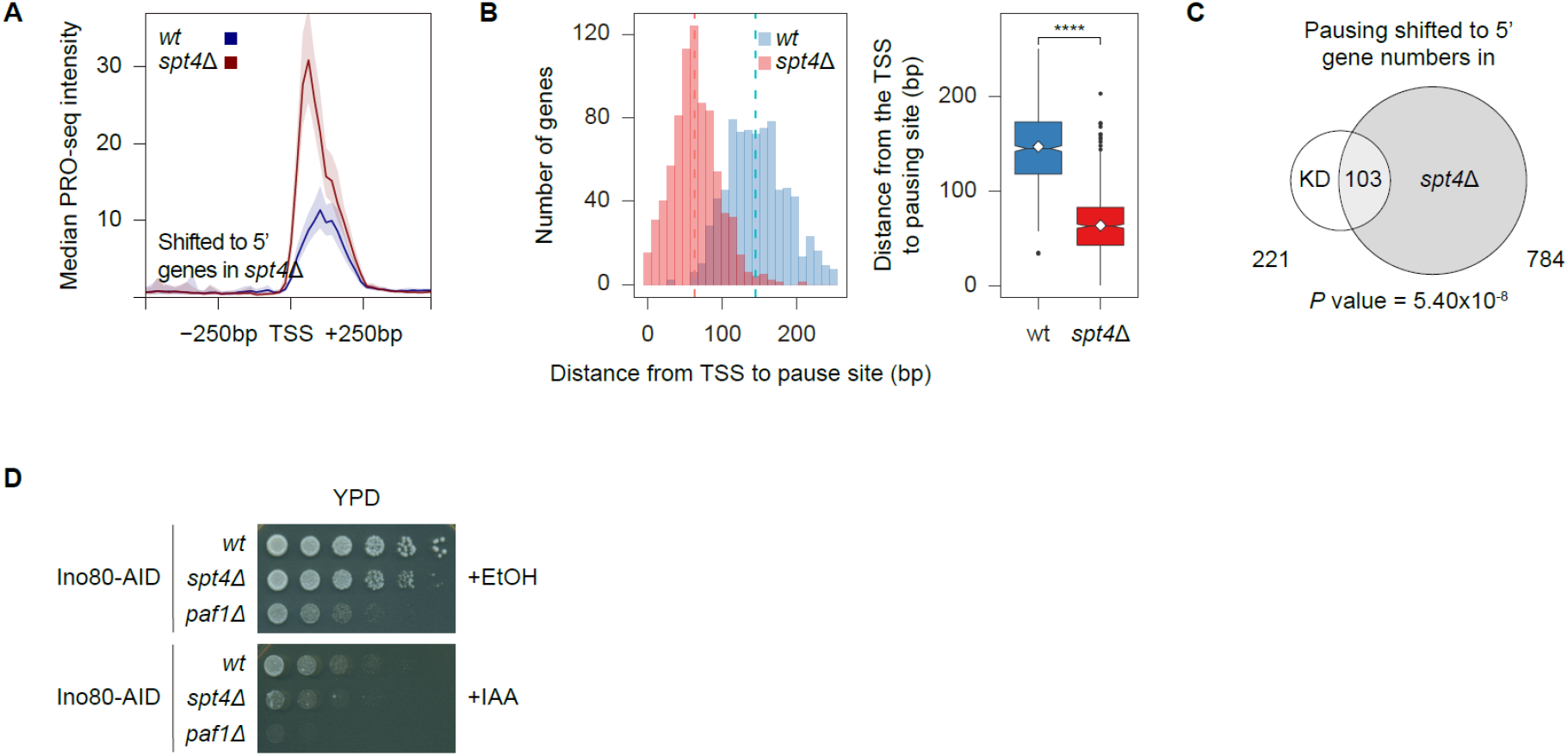
Loss of the conserved pausing factor Spt4p shifts pausing upstream in a manner similar to Ino80p knockdown. (**A**) Average profiles representing median PRO-seq intensities of *wt* and *spt4*Δ. Medians reflect 20-bp bins. (**B**) Histograms and boxplots demonstrating the distance from the TSS to pausing site in *wt* and *spt4*Δ. Asterisks represent statistically significant differences, as calculated using the Wilcoxon test. For (**A**) and (**B**), genes whose pausing sites were shifted toward 5’ (> 30bp) in *spt4*Δ (N = 784) were used for analyses. (**C**) Venn diagram depict overlap between shift-to-5’ genes upon Ino80p-KD (N = 221) and genes showing an upstream shift (> 30bp) of RNAPII pausing in *spt4*Δ (N = 784). *P* value was calculated using the hypergeometric distribution. (**D**) Spotting assays analyzing genetic interactions between *INO80* and *SPT4* or *PAF1*. Cells were spotted onto YPD plates containing either ethanol or auxin (0.5mM) with 5-fold serial dilutions and incubated at 30°C. All PRO-seq data were generated using combined biological replicates. PRO-seq intensities were calculated using spike-in-normalized reads per million.

### The role of Ino80p in RNAPII pausing is conserved in mESCs

Since the Ino80 complex is a highly conserved chromatin remodeler from yeast to humans (*18*), we investigated whether Ino80 loss also caused a defect in elongating RNAPII positioning before pause-release in mESCs. Toward this end, we carried out PRO-seq experiments in mESCs treated with either *siEgfp* or *siIno80* (fig. S4A). To more precisely identify the peak of promoter-proximal pausing, we tiled 1kb around the annotated TSSs (TSS ± 1kb) in a 50-bp window with a 5-bp shift, as previously reported (*4*). We selected the window showing the maximum PRO-seq reads, and designated the 5’ end of the selected window as the “PRO-seq peak”. The average profile of median PRO-seq intensity in mESCs treated with *siEgfp* revealed an almost 2-fold higher PRO-seq intensity at the PRO-seq peak than at the annotated TSS (fig. S4B). Because the PRO-seq signal was highly confined around the PRO-seq peak regions, we determined the regions from 100bp upstream to 200bp downstream of the PRO-seq peak as the promoter-proximal regions for mESCs. Based on the PRO-seq coverage of these regions, we classified genes as being high-confidence paused (N = 27,406; 78.5%) or high-confidence not paused (N = 5,427; 16.3%) among the total protein-coding genes (N = 31,173). To analyze the RNAPII pausing shift under Ino80-KD, we defined the 1^st^ and the 2^nd^ pausing site for each gene in the same manner as in *S. cerevisiae*. To distinguish the pattern of RNAPII transition for each gene, we generated a heatmap of the PRO-seq log_2_ fold change around the pausing site (fig. S4C). However, the results obtained from these analyses differed from those observed in *S. cerevisiae* (fig. S2C and fig. S4C). Although the 2^nd^ pausing site was shifted upstream relative to the 1^st^ pausing site for a small fraction of genes (N = 1,082; 8.9%) upon Ino80-KD, a much larger fraction of genes (N = 2,635; 23.2%) displayed a downstream shift in mESCs (fig. S4C).

Except for the direction of shift, however, Ino80-KD in mESCs yielded a pausing site-determination defect similar to that observed in *S. cerevisiae*. First, the average profile of the median PRO-seq signal around the PRO-seq peak regions for genes whose pausing site shifted more than 30bp in the 3’ direction upon Ino80-KD (“shift-to-3’ genes”; N = 2,324) revealed that Ino80-KD induced prominent downstream movement of RNAPII pausing (Fig. 5A; The arrows and below dotted lines represent the median of the pausing sites). Second, these shift-to-3’ genes exhibited the use of an alternative pausing site rather than a focused pausing peak in the physiological conditions (Fig. 5B). Ino80-KD displaces RNAPII solely at locations where pausing can also occur (Fig. 5B; The 25^th^ and 75^th^ percentiles of the 2^nd^ pausing site relative to the 1^st^ pausing site for shift-to-3’ genes are represented as the two red dotted lines). Third, whereas pausing at the 1^st^ pausing site was decreased by Ino80-KD, pausing at the 2^nd^ pausing site was significantly increased under the same condition (Fig. 5C). The sequence preferences at each pausing site were also similar (fig. S4D). Furthermore, *de novo* motif finding analysis using HOMER (*46*) indicated that the YY1 motif was significantly enriched in the promoter regions of shift-to-3’ genes relative to no-shift genes (fig. S4E). YY1 is physically associated with the mammalian Ino80 complex (*47*), providing additional support for the engagement of Ino80 with these genes. We did not observe a similar motif in *S. cerevisiae*, perhaps reflecting its lack of an identified homolog for YY1 (*48*). Based on our findings, we conclude that Ino80 plays a conserved role in determining where RNAPII should mainly pause at the genes bearing an alternative pausing site in the early transcription elongation stage in mESCs.

**Fig. 5.**
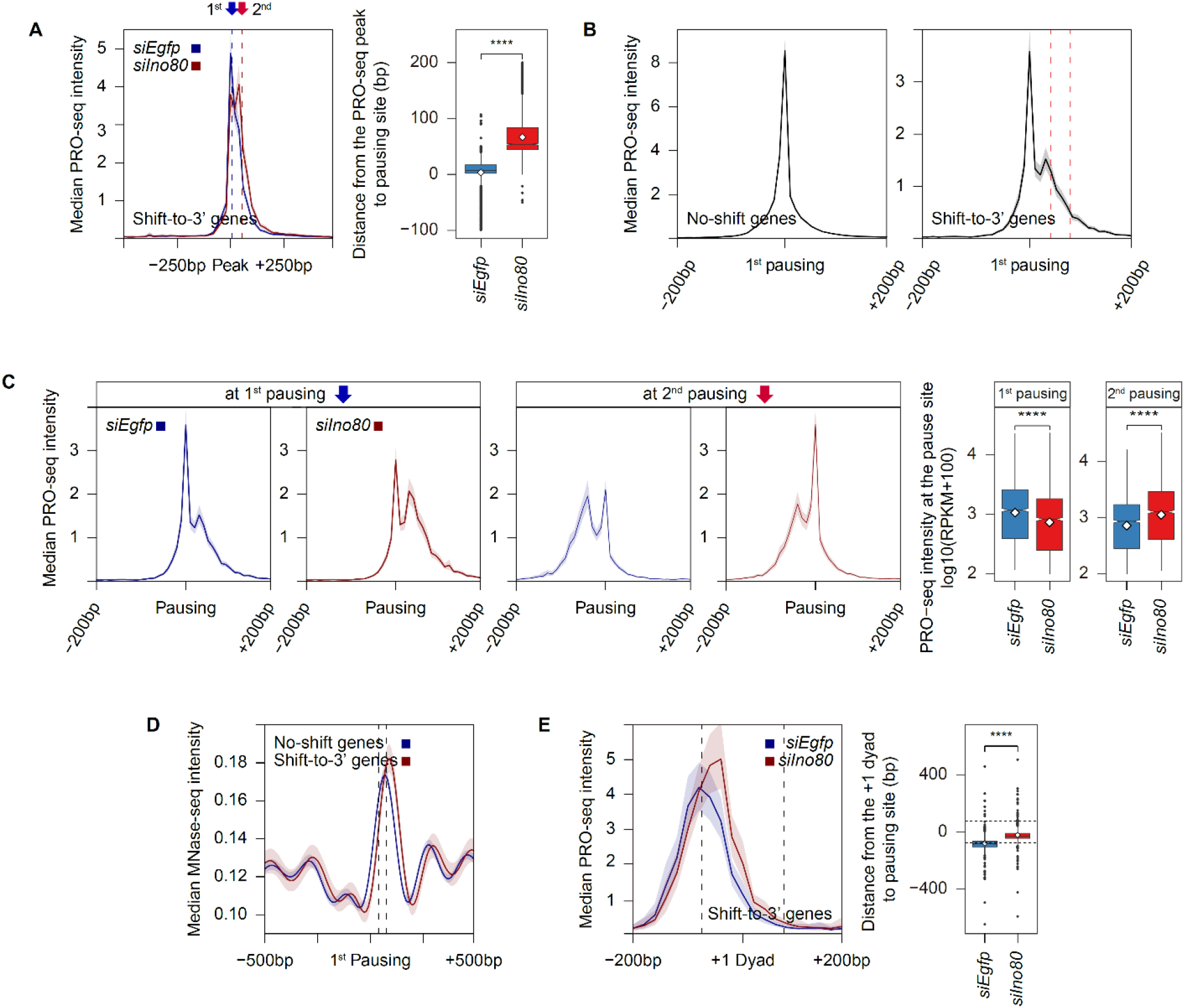
The ability of Ino80 to regulate RNAPII pausing site is conserved in mESCs. (**A**) Average profiles representing median PRO-seq intensities in mESCs treated with either *siEgfp* or *siIno80* for 48 hrs at shift-to-3’ genes (N = 2,324) around the PRO-seq peak. Medians reflect 20-bp bins. The arrows and below dotted lines represent the median of the 1^st^ (blue, 7bp) and 2^nd^ (red, 54bp) pausing sites. Boxplot showed the distance between pausing site and the PRO-seq peak. (**B**) Average profiles of median PRO-seq intensities in mESCs treated with *siEgfp* at shift-to-3’ genes (N = 2,324) or no-shift genes (N = 16,903). Medians reflect 10-bp bins. The two red dotted lines (Right) represent the 25^th^ (39bp) and 75^th^ (76bp) percentiles of the 2^nd^ pausing site relative to the 1^st^ pausing site. (**C**) Average profiles displaying median PRO-seq intensities around the 1^st^ or 2^nd^ pausing site. Medians reflect 10-bp bins. Boxplots showed the PRO-seq intensities at the indicated single nucleotides in log_10_(RPKM+100). (**D**) Average profiles of median MNase-seq intensities obtained from untreated mESCs (GSM2906312 and GSM2906313) for no-shift genes and shift-to-3’ genes at the 1^st^ pausing site. The two black dotted lines indicate the 25^th^ (39bp) and 75^th^ (76bp) percentiles of the 2^nd^ pausing site relative to the 1^st^ pausing site. Medians reflect 10-bp bins. (**E**) Average profiles of median PRO-seq intensities around the +1 dyad, which was determined by MNase-seq used in Fig. 5D, for shift-to-3’ genes. The two dotted lines represent the position of the +1 nucleosome (75 bp upstream and downstream of the +1 dyad). Only nucleosomes overlapped with H3K4me3 enrichment calculated from an existing ChIP-seq data (GSM590111) (*64*) were used, in an effort to exclude false-positive nucleosomes (N = 629). Boxplot indicates the distance from the +1 dyad to pausing site. Asterisks represent statistically significant differences, as calculated using the Wilcoxon test. All PRO-seq data were generated using combined biological replicates. PRO-seq and MNase-seq intensities were calculated using spike-in-normalized reads per million and Gaussian smoothing-normalized reads per million, respectively.

We next deciphered whether the 2^nd^ pausing site was also associated with the +1 nucleosome in mESCs. We analyzed an existing MNase-seq data set obtained from untreated mESCs (*49*). We observed the better positioned +1 nucleosome for shift-to-3’ genes (N = 2,324) relative to no-shift genes (N = 16,903) (Fig. 5D), and found the proximity of the 2^nd^ pausing site to the +1 nucleosome (Fig. 5D; The dotted lines represent the 25^th^ and 75^th^ percentiles of the 2^nd^ pausing site relative to the 1^st^ pausing site). This suggested that the nucleosome context around the 1^st^ pausing site are important for mESCs to suppress the 2^nd^ pausing site, as also found in budding yeast. Consistent with previous reports, the PRO-seq signals corresponding to the 1^st^ pausing site for mammalian shift-to-3’ genes were located near the entrance of the +1 nucleosome (Fig. 5E) (*5, 15, 16*). The PRO-seq signals corresponding to the 2^nd^ pausing site, however, were found between the 1^st^ pausing site and the +1 dyad (Fig. 5E). In control analysis, we detected almost no difference in the PRO-seq pattern around the +1 dyad for no-shift genes (fig. S4F). From these results, we conclude that Ino80 plays a conserved regulatory role in proper localization of RNAPII pausing at the 1^st^ pausing site in a nucleosome-context-dependent manner in various organisms, from budding yeast to mouse (Fig. 6).

**Fig. 6.**
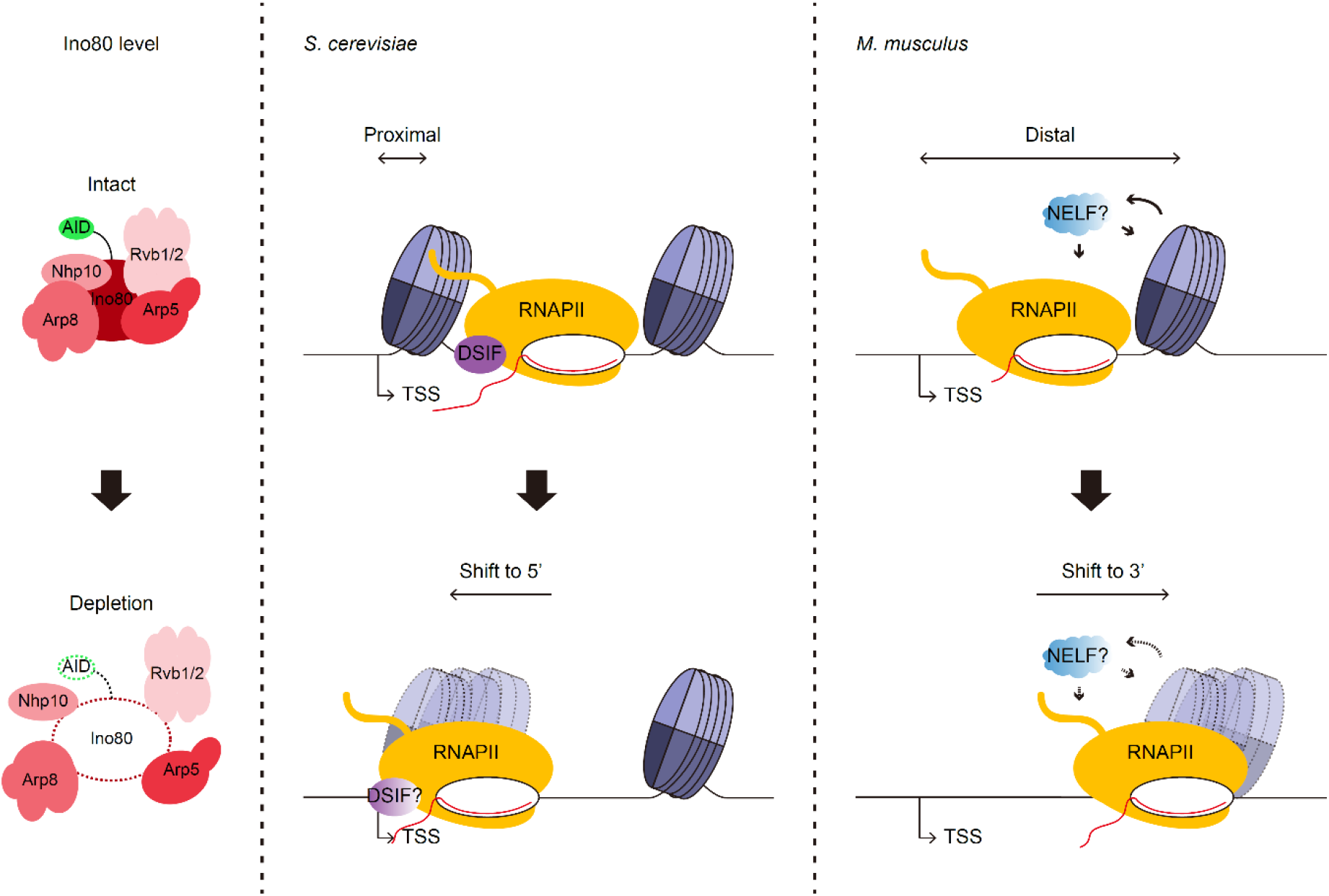
Model of the function of Ino80 in mediating alternative pausing site determination. Model depicts a conserved regulatory role of Ino80 in proper localization of RNAPII pausing through its activity to modulate the +1 nucleosome in budding yeast and mESCs. See Discussion for details.

## Discussion

We found highly confined PRO-seq signals immediately downstream of the observed TSSs in *S. cerevisiae*, which yielded a pattern that was highly similar to that representing promoter-proximal pausing in metazoans. However, the slowed promoter-proximal elongation in *S. cerevisiae* occurred more broadly and RNAPII seemed to be paused farther downstream than in metazoans. Indeed, most of the 1^st^ pausing sites were located downstream of the +1 nucleosome. This finding was unlike that in higher eukaryotes, where most pausing was found to occur upstream of the +1 dyad (*5, 15, 16*). One of the reasons for this discrepancy might be that NELF, which is involved in the pausing of RNAPII at a more promoter-proximal region (*50*), is not conserved in yeasts, including *S. cerevisiae* and *S. pombe* (*51, 52*). The hypothesis is consistent with the PRO-seq distribution reported for *S. pombe*, which showed a near-overlapping association of RNAPII pausing with the +1 dyad (*12*).

Surprisingly, Ino80 knockdown yielded opposite shifts of the RNAPII pausing sites in budding yeast and mESCs. One possible explanation is that Ino80 plays different roles in these species, since the Ino80 complex contains species-distinct components that could target Ino80 to specific contexts. Alternatively, our observation that the nucleosome distribution around the pausing site was similar in the two organisms led us to postulate that these discrepancies could be due to differences in the chromatin architectures around promoter regions in budding yeast versus mESCs (*35, 36*). In *S. cerevisiae*, the +1 nucleosome generally includes a TSS for most genes. Thus, most of the 1^st^ pausing sites are downstream of the +1 nucleosome. The loss of Ino80p induced the main pausing site to shift toward the +1 nucleosome, accompanied by both a moderate increase in nucleosome fuzziness and a decrease in the +1 nucleosome occupancy. This suggests that Ino80p may play a critical role in the passage of elongating RNAPII from the 2^nd^ to 1^st^ pausing site and helps establish intact pausing through its activity to position the +1 nucleosome. Supporting our hypothesis, previous studies showed an increase of nucleosome fuzziness in Ino80p mutants (*21, 41*) and demonstrated that Ino80p functions to pull the +1 nucleosome toward the NFR for a subset of genes (*32*). In higher eukaryotes, the +1 nucleosome is located farther downstream of TSS than in budding yeast. Consistent with this, the median 1^st^ pausing site for Ino80-dependent genes in mESCs was found to be upstream of the +1 nucleosome (Fig. 5E; The median of the *siEgfp* treated samples was 84bp upstream of the +1 dyad). On one hand, this could reflect that Ino80 can promote RNAPII pausing in a nucleosome-independent manner. Alternatively, a previous study implied that the positioned +1 nucleosome recruits more NELF and enhances promoter-proximal pausing (*53*). Thus, mammalian Ino80-dependent nucleosome positioning may also work to establish RNAPII pausing through recruiting *trans*-activating pausing factors. Consistent with this scenario, RNAPII resumed elongation to the 2^nd^ pausing site upon the loss of Ino80 (Fig. 5E). The delocalization of the +1 nucleosome due to Ino80 depletion probably allows further elongation of RNAPII, and the 2^nd^ pausing that occurs around 30bp upstream of the +1 dyad (Fig. 5E; The median of the *siIno80* treated samples) might be due to a physical blocking of the +1 nucleosome. This location seems to be close to the location of SHL(-2), at which transcribing RNAPII can be stalled *in vitro* (*54*). Based on the present and previous findings, we suggested that the nucleosome-dependent intrinsic pausing at the 2^nd^ pausing site is conserved from budding yeast to mESCs, and that Ino80 plays a key role in actively determining the main pausing site through its ability to modulate +1 nucleosome positioning in a context-dependent manner (Fig. 6). In support of our hypothesis, we note that a recent study using human NELF-C-AID suggested that NELF loss results in a similar downstream shift of RNAPII pausing accumulated at the 2^nd^ pausing site, which is tightly associated with the +1 nucleosome (*16*).

Supporting the idea that the between-species discrepancies in RNAPII pausing observed herein are due to differences in chromatin architecture, a defect in DSIF was also reported to cause different phenotype in *S. cerevisiae* versus *M. musculus*. The deletion of Spt4p in budding yeast showed the prominent increase in PRO-seq signal at the 5’ ends of genes (*12*), whereas Spt5 depletion in mouse triggered the accumulation of PRO-seq signals downstream of the promoter-proximal regions (*55*). Interestingly, the directions in which elongating RNAPII accumulated in the above-described mutants are similar to that observed following Ino80-KD in the present work, further supporting that Ino80 and DSIF may play synergistic roles in both species. In addition, Spt4 deletion in *S. pombe* has resulted in PRO-seq distribution resembling a defect upon Spt5 loss in mouse (*12*). Intriguingly, a previous study found a similar but distinct nucleosome distribution in budding and fission yeasts (*56, 57*). Further experiments seeking to gain additional contextual information or identify unknown factors in *S. pombe* might help explain these discrepancies in two divergent yeasts.

Based on our collective results, slowed elongating RNAPII within the promoter-proximal regions is an evolutionarily conserved mechanism from yeast to human. Chromatin remodelers could not only regulate promoter opening and productive elongation but also govern promoter-proximal RNAPII pausing through its role to tune nucleosome architecture. In the future, it will be interesting to study the detailed mechanism of chromatin remodeling in the early transcription elongation.

## Materials and Methods

### Yeast strains and cell culture

The budding yeast strains used in this study are listed in table S2. AID-tagged proteins were conditionally depleted using 250mM auxin (Sigma, I2886) stock dissolved in ethanol at a final concentration of 0.5mM, as previously described (*29*). Briefly, Ino80-AID cells were grown to mid-log phase at 30°C in YPD containing ethanol. The ethanol was removed by media exchange, cells were incubated with auxin (where indicated) for 3 hrs to allow conditional depletion. The auxin was removed by media exchange, and cells were incubated in auxin-free medium an additional 3 hrs. At the indicated time points, Ino80-AID cells were harvested and subjected to PRO-seq and PRO-cap. The workflow is schematically presented in fig. S1C. The efficiency of Ino80p knockdown was confirmed by Western blotting. To generate the deletion strains, we conducted standard LiAc transformation using PCR-based gene targeting. These cells were incubated to mid-log phase at 30°C in YPD and were subjected to PRO-seq.

### mESC culture

E14Tg2a mESCs were maintained under feeder-free conditions at 37°C with 5% CO_2_ in humidified air. Briefly, mESCs were cultured on gelatin-coated dishes in Glasgow’s minimum essential medium (GMEM) containing 10% knockout serum replacement (Gibco, 10828-028), 1% non-essential amino acids (Gibco, 11140-050), 1% sodium pyruvate (Gibco, 11360-070), 0.1 mM β-mercaptoethanol (Gibco, 21985-023), 1% FBS (Hyclone, SH30917.03), 0.5% antibiotic-antimycotic (Thermo, 15140122), and 1,000 units/ml LIF (Millipore, ESG1106).

### RNA interference

The siRNAs against EGFP (5’-GUUCAGCGUGUCCGGCGAG-3’) and Ino80 (5’-GGCUUAUCUGUAAAGGCACAAUUGA-3’) were synthesized and annealed by Bioneer. mESCs were seeded to plates, incubated for 24 hrs, and transfected with the indicated siRNAs (final concentration, 50nM) using DharmaFECT I (Dharmacon, T-2001-03) according to the manufacturer’s protocol. Briefly, 50nM of siRNAs and DharmaFECT I diluted in Opti-MEM (Gibco, 31985062) were separately incubated for 5 min at 25°C, mixed and incubated for 20 min at 25°C, and then used for transfection. The culture medium was replaced at 24 hrs of transfection and cells were harvested at 48 hrs of transfection.

### Western blot analysis of protein depletion

Whole-cell lysates of Ino80-AID cells were prepared using a standard bead-beating protocol, and proteins were eluted by boiling at 100°C for 5 min in 2x SDS sample buffer (20% glycerol, 0.4% bromophenol blue, 100mM Tris-Cl, pH6.8, 4% SDS, and 200mM β-mercaptoethanol). The utilized antibodies included anti-FLAG M2 (Sigma A8592; used at 1:3000) and anti-β-actin (Santa Cruz sc-47778 HRP; used at 1:5000). The AID-tagged Ino80 cells were a gift from the Friedman lab as previously described (*29*). mESCs were washed with PBS (Welgene, LB004-02) and detached from the dishes by incubation with trypsin-EDTA (Gibco, 25300-054) at 37°C for 2 min. The trypsin was inactivated by the addition of GMEM with 1% FBS and 0.5% antibiotic-antimycotic, and the cells were harvested, washed with PBS, and resuspended in EBC300 (120 mM NaCl, 0.5% NP-40, and 50 mM Tris-Cl, pH 8.0) containing protease inhibitors. Whole-cell lysates were prepared by vigorously vortexing the cell mixture for 30 min followed by centrifugation for 30 min at 4°C. Proteins were eluted by being boiled at 100°C for 5 min with 5x SDS sample buffer. The utilized antibodies included anti-Ino80 (Abcam, ab118787; used at 1:1000) and anti-tubulin (Cell Signaling, 2144S; used at 1:4000).

### Spotting assay

Cells were spotted onto the indicated plate in serial five-fold dilutions starting at 0.5OD, and then incubated at 30°C for ∼2 days.

### Yeast cell permeabilization

Yeast cells were permeabilized as described (*6*) with some of previously reported modifications (*58*). Briefly, exponentially growing yeast cells were harvested by centrifugation at 2000rpm for 3min at 4°C. Cells were washed once with ice-cold DEPC-H_2_O. Cell pellets were resuspended in 10ml of 0.5% sarkosyl (Sigma, L5777) and incubated for 20min on ice. Cells were spun down at 400xg for 5min at 4°C and resuspended in storage buffer (10mM Tris-Cl, pH8.0, 25% glycerol, 5mM MgCl_2_, 0.1mM EDTA, and 5mM DTT) to an OD of 5 per 200μl. The solutions were flash-frozen using LN_2_ and stored at −80°C.

### Isolation of nuclei

mESCs were transfected with the indicated siRNAs, and nuclei were isolated as described (*4, 44*) with some modifications. Briefly, ∼20 x 10^6^ plated mESCs were washed once with PBS and detached by incubation with trypsin-EDTA at 37°C for 2min. The trypsin was inactivated by the addition of GMEM with 1% FBS and 0.5% antibiotic-antimycotic, and the cells were harvested and washed twice with ice-cold PBS. The cells were resuspended in 5ml of ice-cold swelling buffer (20mM Tris-Cl, pH7.5, 2mM MgCl_2_, 3mM CaCl_2_, and 2U/ml RNase inhibitor) for 5min on ice. Lysis buffer (5ml; 20mM Tris-Cl, pH7.5, 2mM MgCl_2_, 3mM CaCl_2_, 0.5% NP-40, 10% glycerol, and 2U/ml RNase inhibitor) was added and the cell pellets were resuspended by gentle pipetting using an end-cut tip. The cells were centrifuged at 1000xg for 5min at 4°C and the cell pellets were resuspended in 1ml of freezing buffer (50mM Tris-Cl, pH8.3, 40% glycerol, 5mM MgCl_2_, and 0.1mM EDTA). The pelleted nuclei were transferred into a new 1.5ml tube and were resuspended in freezing buffer at ∼5 x 10^6^ nuclei per 100μl. The solutions were flash-frozen using LN_2_ and stored at −80°C.

### PRO-seq and PRO-cap library preparation

Nuclear run-on reactions and RNA extractions were performed based on the published protocol (*6*) with minor modifications that were previously reported (*44, 58*). Briefly, the flash-frozen yeast cells were quickly thawed on ice. For the yeast spike-in control, 0.125OD of permeabilized *S. pombe* (ED665) cells were added to each 5OD of permeabilized *S. cerevisiae* sample before the nuclear run-on reaction was performed. Combined yeast cells were spun down at 400xg for 5min at 4°C. Nuclear run-on reactions were conducted with 25μM biotin-11-UTP (PerkinElmer, NEL543001EA), 25μM biotin-11-CTP (PerkinElmer, NEL542001EA), 125μM ATP (Roche, 11140965001), and 125μM GTP (Roche, 11140957001) in run-on reaction buffer (20mM Tris-Cl, pH7.7, 200mM KCl, 5mM MgCl_2_, 2mM DTT, and 0.4U/μl RNase inhibitor) with 0.5% sarkosyl. The reaction mixtures were incubated at 30°C for 5min. For the isolated nuclei of mESCs, nuclear run-on reactions were performed with 25μM biotin-11-UTP (PerkinElmer, NEL543001EA), 25μM biotin-11-CTP (PerkinElmer, NEL542001EA), 125μM ATP (Roche, 11140965001), and 125μM GTP (Roche, 11140957001) in run-on reaction buffer (5mM Tris-Cl, pH8.0, 150mM KCl, 2.5mM MgCl_2_, 0.5mM DTT, and 0.4U/μl RNase inhibitor) with 0.5% sarkosyl. The reaction mixtures were incubated at 37°C for 5 min.

RNA was extracted from the run-on-reacted cell pellets using either a standard hot-phenol method (for yeast samples) or TRIzol LS (Ambion, 10296028; for mESC samples). Next, the respective library was generated followed using the published PRO-seq or PRO-cap protocols (*6*) for the steps spanning RNA fragmentation by base hydrolysis to full-scale PCR amplification. Note that there were a few differences in the applied reagents: we used Superscript IV reverse transcriptase (Invitrogen, 18091050) instead of Superscript III (Invitrogen, 56575); we used 25mM of each dNTP (Thermo Scientific, R1121) instead 12.5mM of each dNTP (Roche, 03622614001); and we used Phusion High-Fidelity DNA Polymerase (Thermo Scientific, F530L) instead of Phusion polymerase (NEB, M0530). DNA libraries of ∼140 bp to 350 bp were selected by agarose gel extraction (Zymo Research, D4007) according to the manufacturer’s protocol and sequenced using an Illumina HiSeq X Ten, HiSeq 4000 and NovaSeq 6000.

### Sequence alignment and data processing (PRO-seq and PRO-cap)

Sequence alignment and data processing were performed based on the publicly available alignment pipelines of GitHub, as used in the previous study (*58*) with minor modifications. Briefly, raw sequencing reads were processed using FASTX-Toolkit (http://hannonlab.cshl.edu/fastx_toolkit/) as follows: Adaptor sequences (5’-TGGAATTCTCGGGTGCCAAGG-3’) were removed, the reads were trimmed to a maximum length of 36nt and, for PRO-seq, the reads were reverse-complemented. Next, reads that mapped to rRNA sequences were depleted using SortMeRNA (*59*), and reads that were not mapped to rRNA sequences were uniquely aligned to the genome using Bowtie, with the algorithm allowing for two mismatches (*60*): The processed reads of yeast samples that were generated with the spike-in approach were mapped to a combined genome consisting of *S. cerevisiae* (sacCer3) and *S. pombe* (SpombeASMv2), and then uniquely aligned reads from each genome were parsed for downstream analysis. The processed reads of mESCs samples were mapped to the *M. musculus* mm10 genome. The coverage of the aligned reads was generated using the genomecov function of BEDtools (*61*). Only the most 3’ nucleotide of each read was calculated for PRO-seq, and only the most 5’ nucleotide of each read was calculated for PRO-cap. For the spike-in control, the recorded coverage in the bedGraph file was normalized by the relative number of reads mapped to a *S. pombe* genome (table S1). bedGraph files were converted to BigWig files by bedGraphToBigWig (*62*) and the downstream analysis was performed based on the publicly available custom R scripts on GitHub, as previously reported (*58*).

Protein-coding gene sets based on the annotated data in the Saccharomyces Genome Database (SGD; N = 6,692) were initially used for *S. cerevisiae* samples. The observed TSS was defined as the single nucleotide with the most PRO-cap reads within the 250bp upstream and downstream of the annotated TSS, in a similar manner to that used in the previous report (*12*). Genes with no PRO-cap signal, genes that had PRO-seq reads lower than 10, and genes shorter than 300bp were filtered out; in the end, 5,715 genes were used out of 6,692 SGD genes. For mESCs, protein-coding gene sets based on the RefSeq annotation were downloaded from UCSC Genome Browser. Genes that had PRO-seq reads lower than 10 and those shorter than 1kb were discarded; in the end, 31,173 genes were selected out of 37,802 genes. The annotated TSS ± 1kb was tiled in a 50-bp window by shifting 5 bp, and the PRO-seq coverage for each window was measured. The window of the most PRO-seq reads was selected and the 5’ position of the selected window was used as the PRO-seq peak, in a manner similar to that described in a previous study (*4*). To identify paused gene sets, mappable reads within the promoter-proximal regions and gene-body regions were calculated. Significant enrichments of the signals from the promoter-proximal regions compared to those from the gene-body regions were evaluated using Fisher’s exact test with Bonferroni’s correction. For *S. cerevisiae*, the regions from the observed TSS to downstream 250bp (TSS to TSS+250 bp) were used as the promoter-proximal regions and those from TSS + 250bp to the annotated transcription end site (TES) were used to calculate the gene-body signals. For mESCs, the regions from 100bp downstream to 200bp upstream of the observed PRO-seq peaks were used as the promoter-proximal regions, and gene-body signal was measured from the regions of 1kb downstream of the observed PRO-seq peak to the annotated TES. A gene was identified as being paused if the *P* value was lower than 0.01 and as being not paused if the *P* value was higher than 0.99, as determined using combined replicates. A gene exhibiting *P* value < 0.05 or *P* value > 0.95 for both biological replicates was further assigned as a high-confidence paused or high-confidence not paused gene, respectively. Pausing site was defined as the single-nucleotide of the maximum PRO-seq read within the promoter-proximal regions, as described in a recent study (*16*). If multiple sites with the maximum values were detected, the closest one to the TSS was selected.

All average profiles centered on the indicated point in this work were generated using a bootstrapped estimation. Briefly, 1,000 random gene sets were taken as each representing 10% of the total genes, and the median and confidence intervals of each averaged subsample were calculated. In the relevant figures, the thick line represents the median value and shaded regions indicate the 12.5^th^ and 87.5^th^ percentiles.

### Analysis of publicly distributed ChIP-seq and MNase-seq data

Raw sequencing reads of the indicated accession numbers were downloaded from NCBI GEO, unless otherwise noted. For MNase-seq, raw reads were uniquely mapped to the *S. cerevisiae* sacCer3 genome or to the *M. musculus* mm10 genome using Bowtie, which trimmed the 3’ bases to 36 bp (if the raw reads were longer than 36 bp), allowed two mismatches, and restricted the maximum insert size to 200bp. BEDtools was used to covert the aligned BAM files to BED formats. The BED files were then processed by iNPS (*63*) to determine the nucleosome positions. Briefly, the “MainPeak” nucleosome that was the closest to either the observed TSS in *S. cerevisiae* or the observed PRO-seq peak in mESCs was assigned as the +1 nucleosome. The +1 dyad was defined as the mid-point between the start and the end inflection, and 75 bp around the +1 dyad was referred to as the +1 nucleosome position. To discard false-positive nucleosome positions, nucleosomes that did not overlap the H3K4me3 ChIP-seq enrichment calculated from the existing data (*34, 64*) were discarded, as previously reported (*16*). The Gaussian smoothing values were used to process BigWig files for average profiles. For ChIP-seq, a combined genome consisting of *S. cerevisiae* (sacCer3) and *S. pombe* (SpombeASMv2) was used, and unique reads from each genome were parsed for downstream analysis. MACS2 (*65*) was used to convert the aligned BAM files to bedGraph formats. For the spike-in control, the recorded coverage in the bedGraph file was normalized by the relative number of reads mapped to a *S. pombe* genome. The bedGraph files were converted to BigWig files by bedGraphToBigWig (*62*).

### Statistical analysis

Statistical analyses were performed using R 3.6.3. In boxplots, whiskers represent 1.5 x interquartile range. *P* values for boxplot were calculated using stat_compare_means (method = “wilcox”) function in ggpubr library. Symbols of ns, *, **, ***, **** represent *P* > 0.05, <= 0.05, <= 0.01, <= 0.001, <= 0.0001 respectively. *P* values for Venn diagram were calculated by hypergeometric distribution using phyper function in stats library.

## Acknowledgments

**General** We would like to thank Nir Friedman for providing the AID strain. We acknowledge Gregory T. Booth and John T. Lis for making publicly available the custom script for PRO-seq analysis in GitHub. We also thank the Lee lab members, H. Jo for obtaining the AID strain from N. Friedman, H. Yang for assistance with mESC culture and Y. Chun for critically reading the paper. **Funding:** This work was supported by a National Research Foundation (NRF) of Korea Grant funded by the Ministry of Science and ICT (MSIT) (2018R1A5A1024261, SRC), and the Collaborative Genome Program for Fostering New Post-Genome Industry of the NRF funded by the MSIT (2018M3C9A6065070). **Author Contributions:** Y.C. and D.L. conceived the study and designed the experiments. Y.C. performed all the PRO-seq experiments and computational analysis in this study. Y.C. and D.L. wrote the paper. S.H. conducted the mESC cultures and siRNA transfections. T.K. supported the computational analysis. **Competing interests:** We have no conflicts of interest to disclose.

## Supplementary Materials

**fig. S1.**
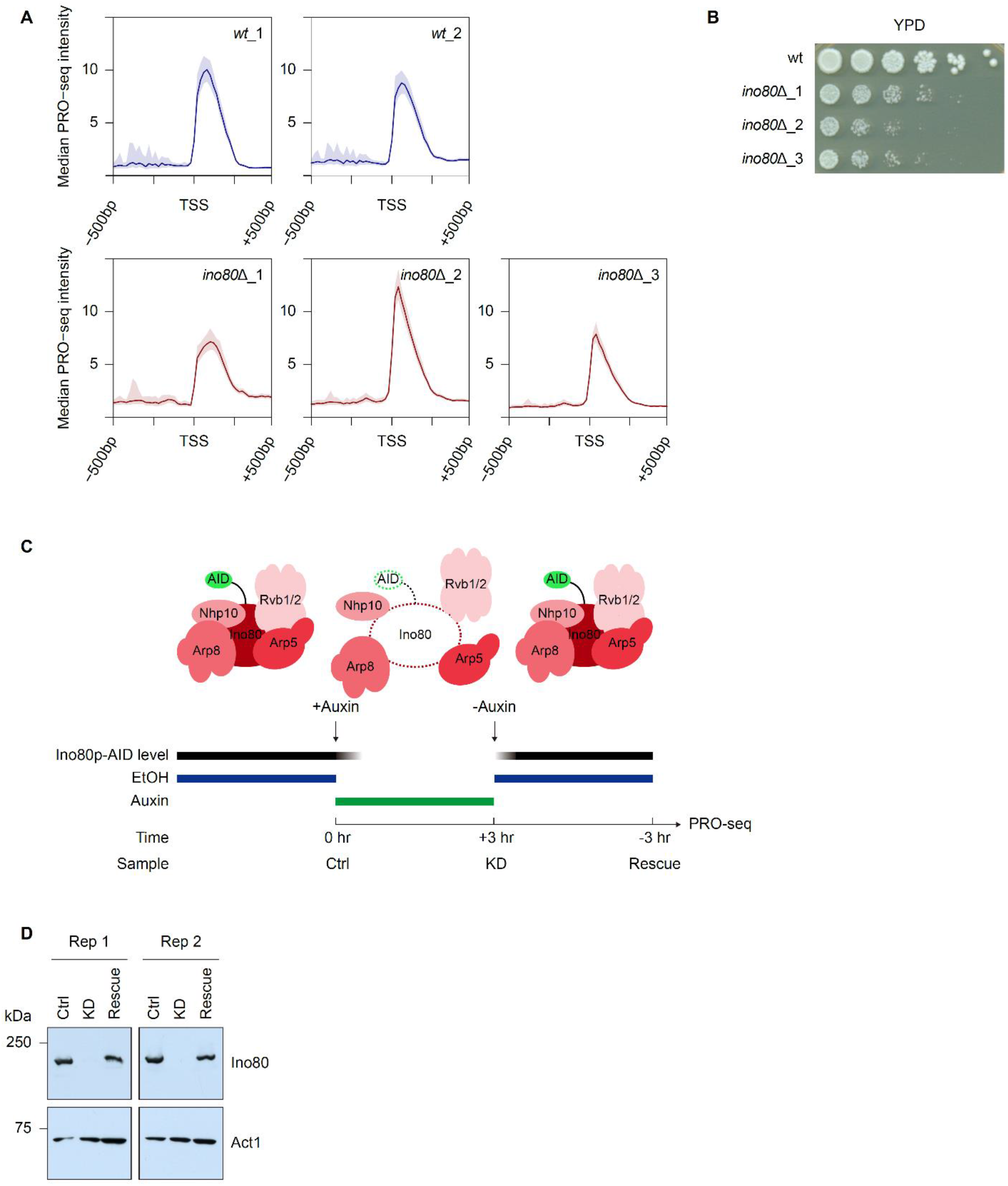
Loss of Ino80p causes variation in fitness and the promoter-proximal PRO-seq pattern. (**A**) Average profiles indicating the median PRO-seq intensities (Spike-in-normalized reads per million) in *wt* and *ino80*Δ. Medians reflect 20-bp bins. (**B**) Spotting assays assessing growth variation in *ino80*Δ cells of the same batch used to generate the triplicate PRO-seq data presented in fig. S1A. Cells were spotted onto YPD plates with 5-fold serial dilutions and incubated at 30°C. (**C**) Schematic illustration of experimental outline. Ino80-AID cells (*29*) were grown to mid-log phase in YPD containing ethanol (Ctrl). The ethanol was washed away and the cells were incubated with auxin (0.5mM) for 3 hrs (KD). The auxin was washed away and the cells were incubated without auxin for additional 3 hrs (Rescue). PRO-seq was performed at each indicated time point. (**D**) Western blot of whole-cell lysates from Ino80-AID cells under the Ctrl, KD, and Rescue conditions showed that Ino80p is almost completely degraded after 3 hrs of auxin incubation and restored after 3 hrs of auxin withdrawal. Western blot analysis was performed against Act1p served as a loading control.

**fig. S2.**
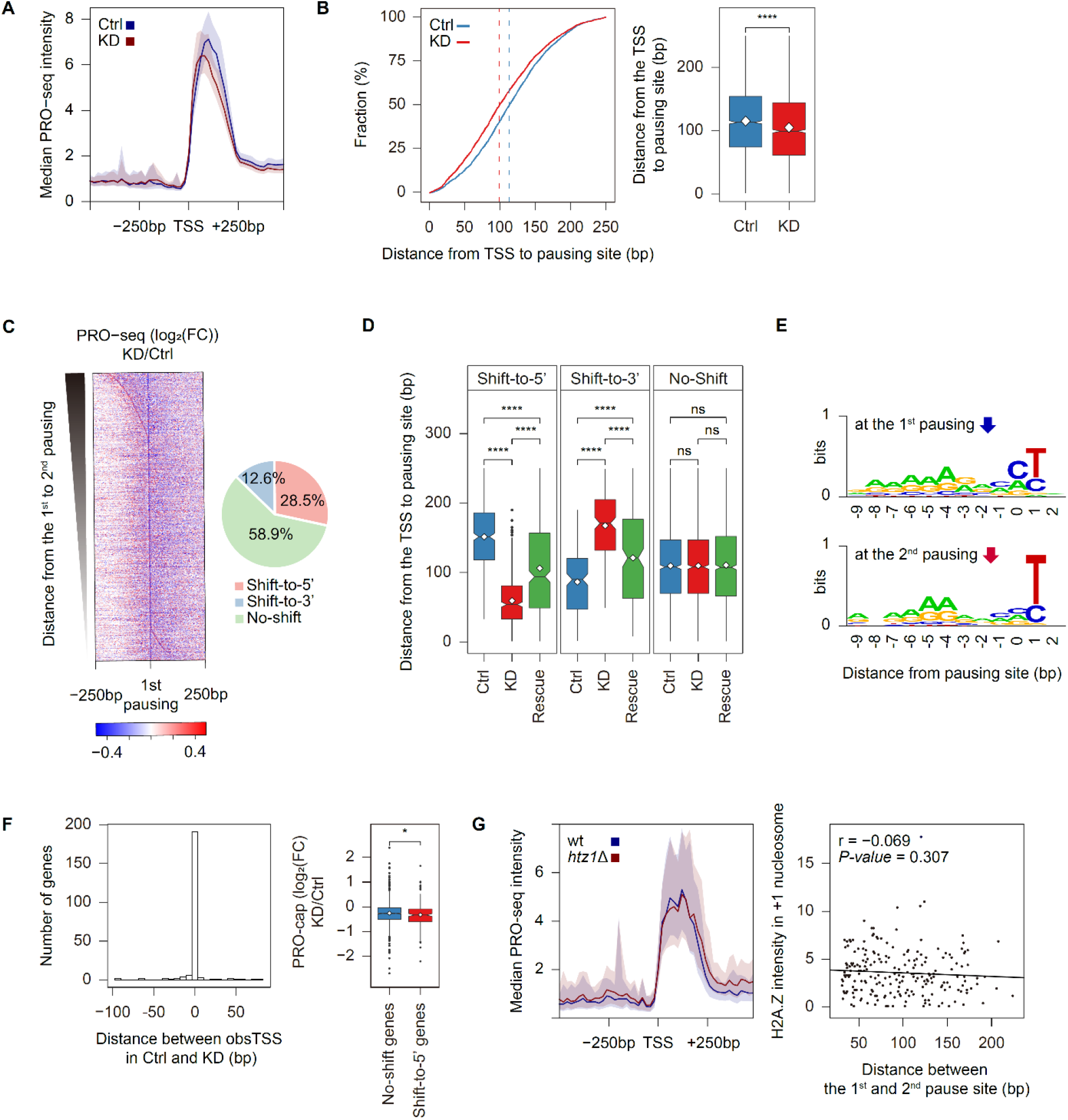
Ino80p knockdown transits the RNAPII pausing from the 1^st^ to the 2^nd^ pausing site and it is independent of both TSS usage and H2A.Z^Htz1^. (**A**) Average profile indicating median PRO-seq intensities in Ino80-AID cells treated with auxin for 0 hr (Ctrl) and 3 hrs (KD), as assessed at the genes defined as being high-confidence paused (N = 2,599). (**B**) Cumulative curve and boxplot demonstrate the distance from the TSS to pausing site for paused genes in the control and knockdown conditions. The dotted lines represent the median value (113bp for Ctrl and 99bp for KD). (**C**) Heatmap representing the PRO-seq signal upon Ino80p-KD as a log_2_ fold change relative to the control sample at the 1^st^ pausing site. Genes were sorted by the distance to the 2^nd^ pausing site. Pie chart shows the percentage of genes whose pausing site was shifted toward 5’ (N = 742), toward 3’ (N = 327), or not shifted (N = 1,530) among total paused genes (N = 2,599) upon Ino80p-KD. (**D**) Boxplot indicating the distance from the TSS to pausing site for genes shifted more than 30bp toward upstream (N = 442) or downstream (N = 178), or for not shifted genes (N = 1,530). Upon rescue of Ino80p, the pausing site was restored to the 1^st^ pausing site in both types of shifted genes. (**E**) Sequence logos around either the 1^st^ or the 2^nd^ pausing site were generated using WebLogo (*37*). (**F**) Histogram analyzing the distance between the observed TSS for shift-to-5’ genes under control and Ino80p-KD conditions (Left). A large fraction of genes (189 out of 221) showed no change in their major initiation site upon knockdown. Boxplot demonstrates the log_2_ fold change in the PRO-cap signal 100bp around the TSS (in RPKM) upon Ino80p-KD relative to control for shift-to-5’ genes or no-shift genes (Right). (**G**) Average profiles indicating the median PRO-seq intensities in *wt* (BY4741) and *htz1*Δ cells (Left). Scatter plot displays the correlation (as Pearson’s r) between the H2A.Z^Htz1^ intensity (IP/input in reads per million) at the +1 nucleosome and the distance of the pausing site shift (Right). For analysis of H2A.Z intensity, we used an existing MNase-ChIP-seq data set (GSM2790633, GSM2790634, GSM2790635 and GSM2790636). All PRO-seq data were generated using combined biological replicates. The PRO-seq and PRO-cap intensities were calculated using spike-in-normalized reads per million. For average profiles, medians reflect 20-bp bins. Asterisks represent statistically significant differences, as calculated using the Wilcoxon test.

**fig. S3.**
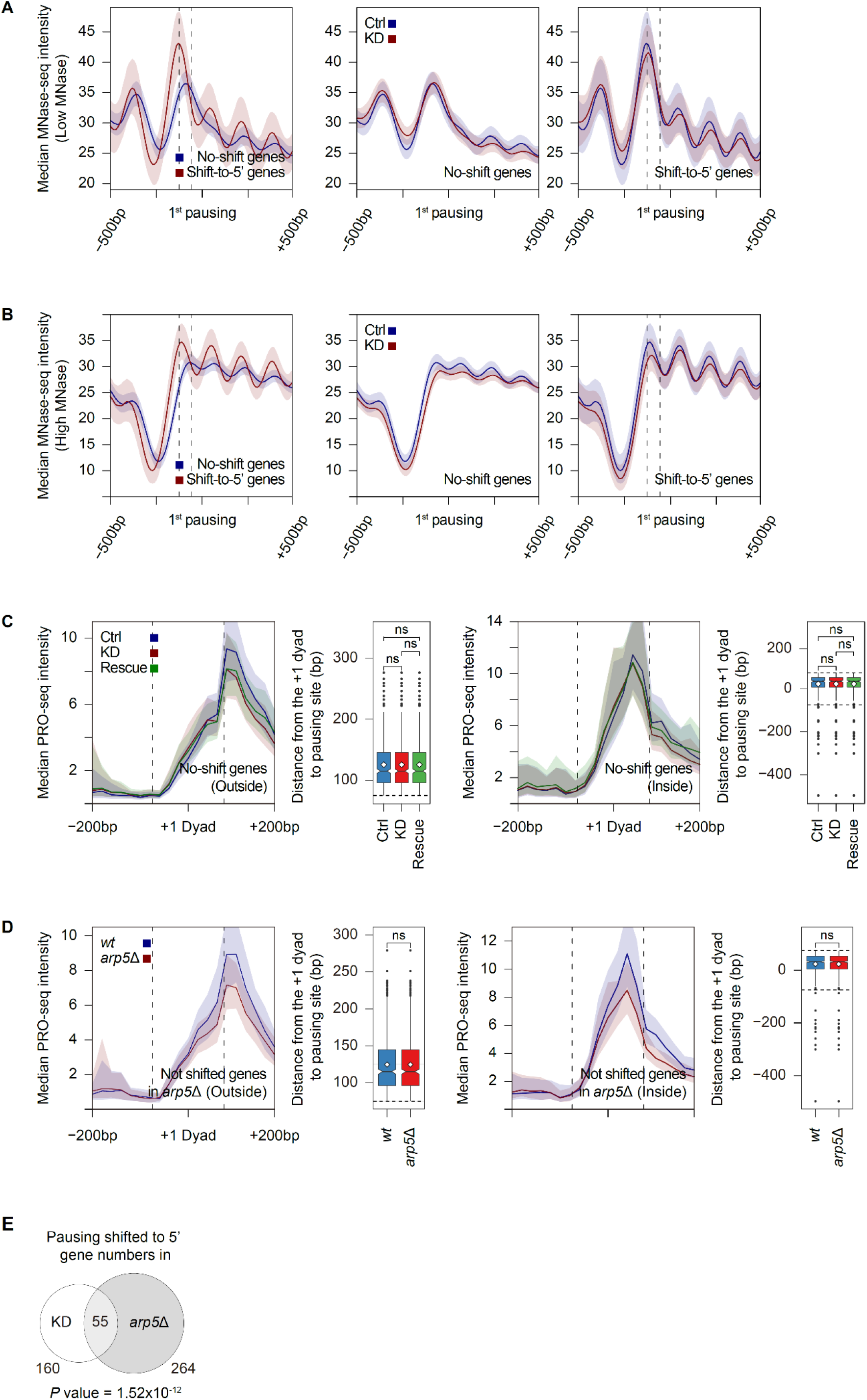
MNase-seq analyses at the pausing site and PRO-seq analyses at the +1 dyad. (**A**) Average profiles of median MNase-seq intensities in control samples (GSM3177776) for no-shift genes (N = 1,211) and shift-to-5’ genes (N = 221) centered on the 1^st^ pausing site (Left). Average profiles of median MNase-seq intensities in control sample (GSM3177776) and Ino80p-KD sample (GSM3177778) for no-shift genes (Middle) and shift-to-5’ genes (Right) at the 1^st^ pausing site. (**B**) Same as (**A**). Instead, GSM3177777 was used as control samples and GSM3177779 was used as Ino80p-KD samples. For fig. S3, A and B, the two dotted lines indicate the 25th (-56 bp) and 75th (-126 bp) percentiles of the 2^nd^ pausing site relative to the 1^st^ pausing site. (**C** and **D**) Average profiles of median PRO-seq intensities for the indicated samples around the +1 dyad (defined in Fig. 1F) and boxplots indicate the distance from the +1 dyad to pausing site. Note that only nucleosomes overlapped with H3K4me3 ChIP-seq enrichment (GSM2507874) were assigned to the +1 nucleosome, in an effort to exclude false-positive nucleosomes. No-shift genes upon Ino80p-KD for (**C**) and not shifted genes in *arp5*Δ for (**D**) were divided into two groups, namely those exhibiting a pausing site either outside (N = 463 for Ino80p-KD and N = 518 for *arp5*Δ; Left) or inside (N = 401 for Ino80p-KD and N = 603 for *arp5*Δ; Right) of the +1 nucleosome, which allowed us to more clearly distinguish the location of the PRO-seq peak. The two dotted lines represent the position of the +1 nucleosome (75 bp upstream and downstream of the +1 dyad). Asterisks represent statistically significant differences, as calculated using the Wilcoxon test. (**E**) Overlap between shift-to-5’ genes upon Ino80p-KD (N = 160) and genes showing an upstream shift (> 30bp) of RNAPII pausing in *arp5*Δ (N = 264). *P* value was calculated using the hypergeometric distribution. All PRO-seq data were generated using combined biological replicates. PRO-seq and MNase-seq intensities were calculated using either spike-in-normalized reads per million or gaussian smoothing-normalized reads per million, respectively. For average profiles, medians reflect either a 20-bp bin (PRO-seq) or a 10-bp bin (MNase-seq).

**fig. S4.**
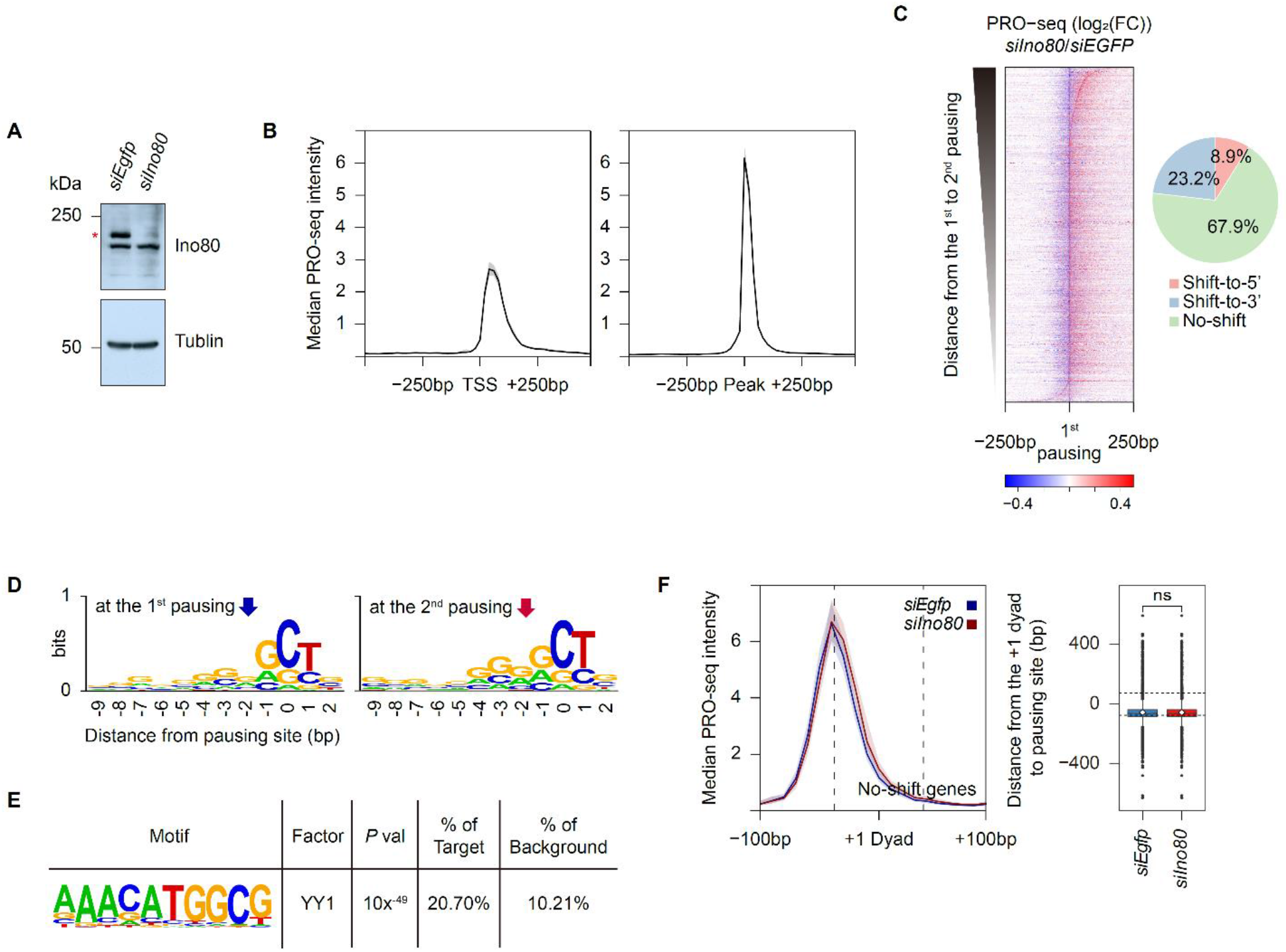
Ino80 knockdown yields a conserved defect of RNAPII pausing in mESCs. (**A**) Western blot analysis of whole-cell lysates from mESCs treated with *siEgfp* or *siIno80* for 48 hrs shows that Ino80 was almost completely and specifically degraded in the presence of *siIno80* (The red asterisk). Tubulin was detected as a loading control. (**B**) Average profile of median PRO-seq intensities in mESCs treated with *siEgfp* centered on either the annotated TSS or the observed PRO-seq peak. (**C**) Heatmaps representing PRO-seq signals at the 1^st^ pausing site in mESCs treated with *siIno80*, given as a log_2_ fold change relative to that in mESCs treated with *siEgfp*. Genes were sorted by the distance to the 2^nd^ pausing site. Pie chart shows the percentage of genes whose pausing sites are shifted toward 5’ (N = 2,206), toward 3’ (N = 5,767), or not shifted (N = 16,903). (**D**) Sequence logos around either the 1^st^ or 2^nd^ pausing site were generated using WebLogo (*37*). (**E**) *De novo* Motif analysis using the findMotifsGenome.pl program of HOMER (*46*) found that the YY1 motif was significantly enriched at the promoter regions of shift-to-3’ genes (N = 2,324) compared to no-shift genes (N = 16,903). The sequences 250bp upstream and downstream of the observed PRO-seq peak were used as the promoter regions. (**F**) Average profiles of the median PRO-seq intensity around the +1 dyad, which was determined by MNase-seq used in Fig. 5D, for no-shift genes. The two dotted lines represent the positions of the +1 nucleosome (75 bp upstream and downstream of the +1 dyad). Note that only nucleosomes with H3K4me3 ChIP-seq enrichment (GSM590111) were used, in an effort to exclude false-positive nucleosomes (N = 4,564). Boxplot indicates the distance from the +1 dyad to pausing site. All PRO-seq data were generated using combined biological replicates. PRO-seq intensity was calculated in reads per million. For average profiles, medians reflect 20-bp. For heatmaps, the signals reflect a 10-bp bin around the indicated site. Asterisks represent statistically significant differences, as calculated using the Wilcoxon test.

**table S1.**
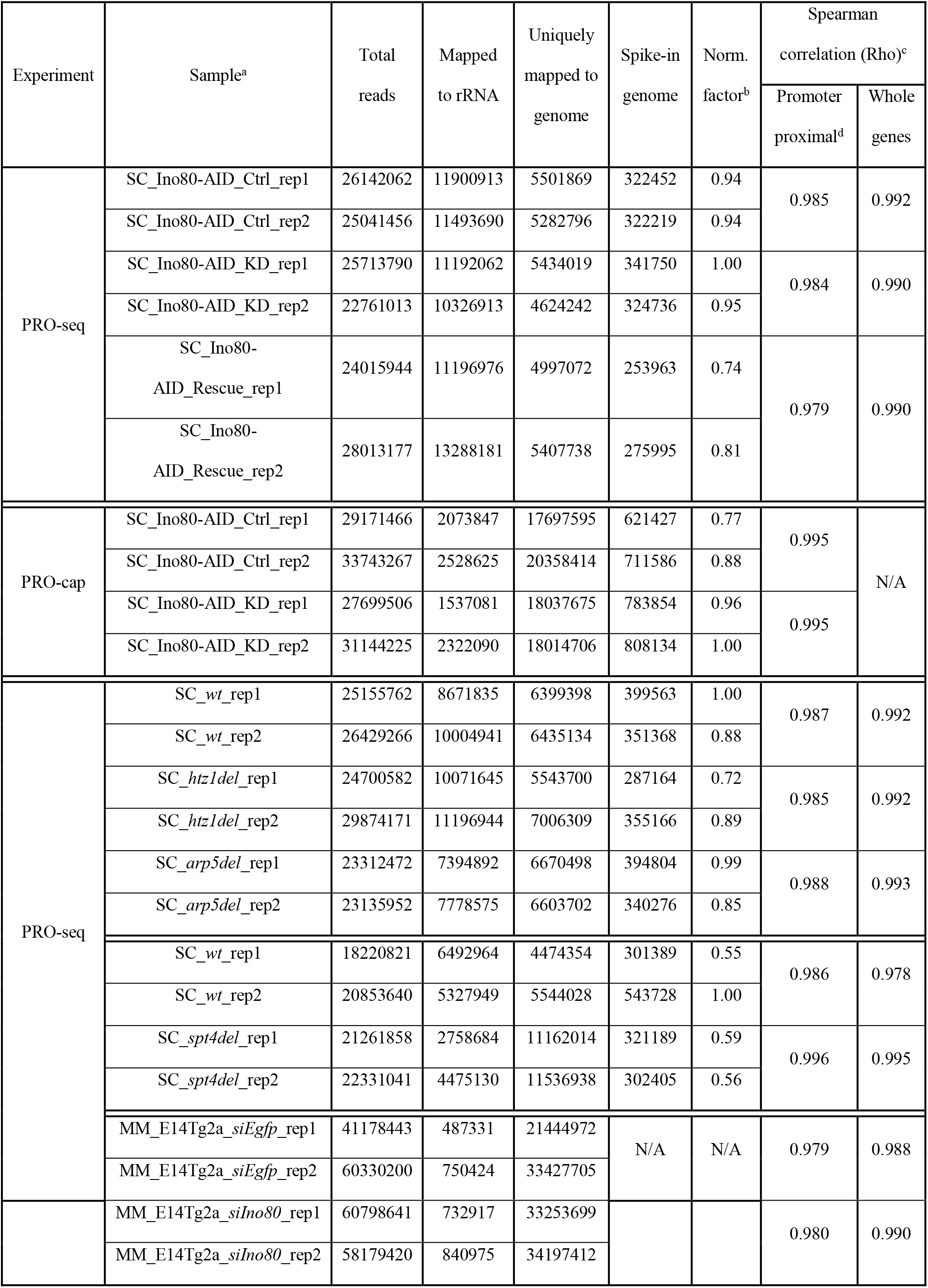
Summary of PRO-seq reads and reproducibility obtained in this study. ^a^ SC indicates *S. cerevisiae* sample and MM indicates mESCs sample. ^b^ Norm. factor was calculated by the relative number of reads mapped to a *S. pombe* genome. ^c^ Reproduciblity of PRO-seq and PRO-cap was calculated by a Spearmans’ Rho using spike-in-normalized reads per million at the indicated regions. ^d^ For *S. cerevisiae*, the regions from the observed TSS to downstream 250bp (TSS to TSS+250 bp) were used as the promoter-proximal regions. For mESCs, the regions from 100bp downstream to 200bp upstream of the observed PRO-seq peaks were used as the promoter-proximal regions.

**table S2.**
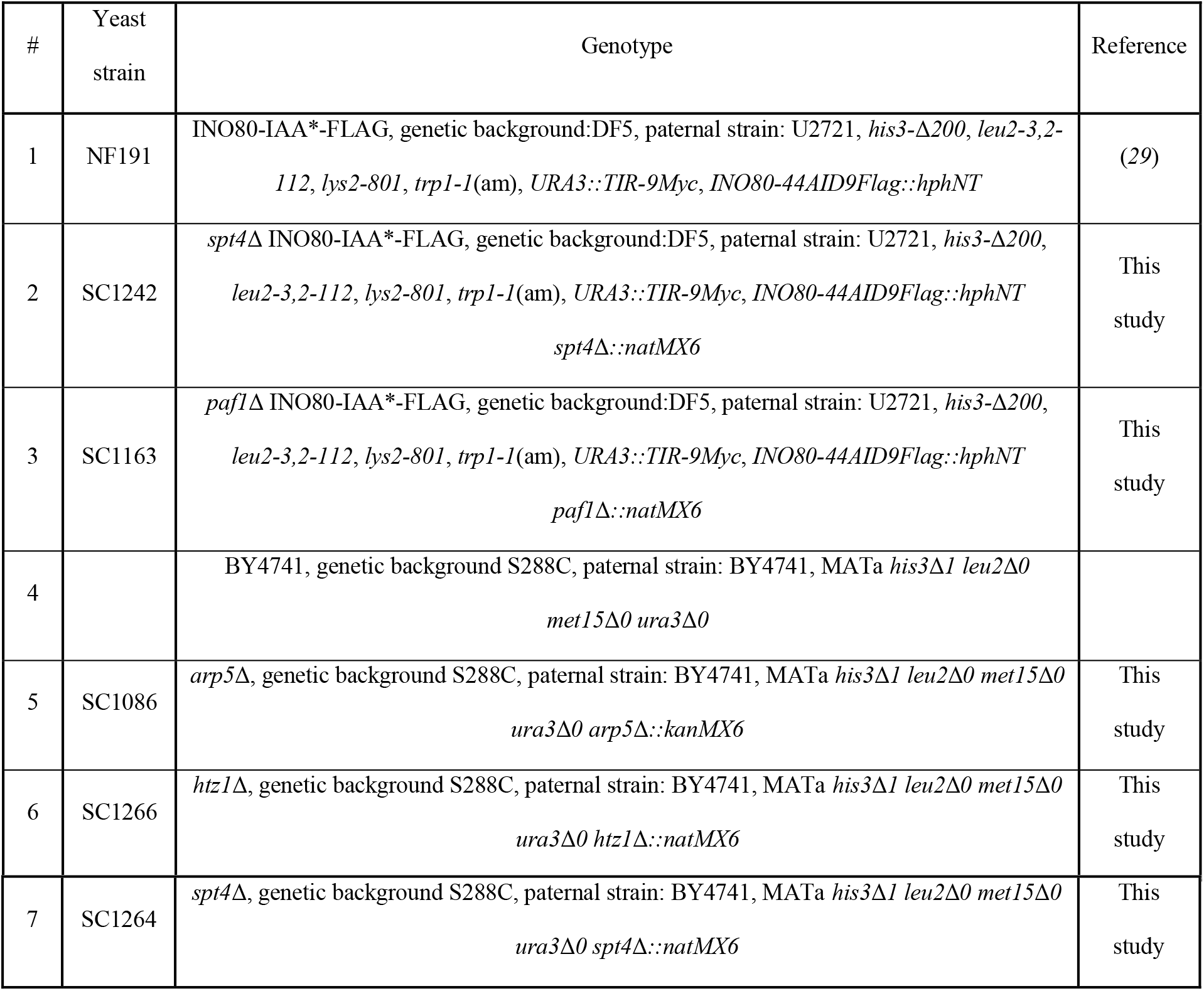
List of strains used in this study.

## Notes

### Competing Interest Statement

The authors have declared no competing interest.

